# Genomic, Clinical, and Spatial Predictors of Durable Response to BRAF/MEK Inhibition in *BRAF*-Mutant Melanoma

**DOI:** 10.64898/2026.05.09.721157

**Authors:** Yingxiao Shi, Alexander Savchenko, Jan Christoph Brase, Brendan Reardon, Cora A. Ricker, Jihye Park, Giuseppe Tarantino, Michael P. Manos, Amy Y. Huang, Eliezer M. Van Allen, Levi A. Garraway, Keith T. Flaherty, David Liu

**Affiliations:** Division of Population Sciences, Dana-Farber Cancer Institute, Boston, MA, USA; Broad Institute of MIT and Harvard, Cambridge, MA, USA; Laboratory of Systems Pharmacology, Harvard Program in Therapeutic Science, Harvard Medical School, Boston, MA, USA; Novartis, Cambridge, MA, USA; Novartis Pharma AG, Basel, Switzerland; Department of Systems Biology, Harvard Medical School, Boston, MA, USA; Parker Institute for Cancer Immunotherapy, Dana-Farber Cancer Institute, Boston, MA, USA; Department of Medical Oncology, Dana-Farber Cancer Institute, Boston, MA, USA; Harvard Medical School, Boston, MA, USA; Computational and Systems Biology Program, Massachusetts Institute of Technology, Cambridge, MA, USA; Massachusetts General Hospital, Boston, MA, USA; Roche/Genentech, South San Francisco, CA, USA

## Abstract

BRAF-targeted therapy (BRAFi/MEKi) and immune checkpoint blockade (anti–PD-1/anti-CTLA-4) have transformed the treatment of *BRAF*-mutant metastatic melanoma. While most patients who respond to targeted therapy eventually progress, a subset derives durable benefit, and biomarkers to identify this subset would inform optimal treatment selection. In this study, we analyzed pre-treatment tumor samples from a clinically annotated cohort of 155 patients with *BRAF*-mutant metastatic melanoma treated with first-line BRAFi/MEKi and followed for up to five years. We stratified patients into durable responders (PFS ≥ 24 months) and rapid progressors (PFS < 6 months with progression) and found that a global metric of tumor genomic heterogeneity, rather than individual gene alterations, distinguished these groups. Combining genomic heterogeneity with baseline tumor burden (e.g., lactate dehydrogenase (LDH) or radiographic lesion dimensions), we developed a parsimonious model that predicted durable responders with high precision and specificity. Notably, the analogous population of patients treated instead with immunotherapy were not durable responders, suggesting that the selected predictors of durable responders are targeted therapy specific. Spatial profiling of a subset of pre-treatment biopsies (n = 47) demonstrated that high intratumoral, but not peritumoral, CD8^+^ T-cell infiltration correlated with prolonged survival on BRAF-targeted therapy and served as an independent predictive factor when considered with genomic heterogeneity and features of clinical tumor burden. Together, these findings highlight the distinct baseline intrinsic and extrinsic features underlying durable response to BRAF-targeted therapy and support their potential implication in guiding treatment selection for patients with *BRAF*-mutant metastatic melanoma.

**One-Sentence Summary:** Integrated clinical, tumor genomic, and immune microenvironmental features predict durable responses to BRAF-targeted therapy.

## INTRODUCTION

The treatment landscape for metastatic melanoma has been transformed, driven by an improved understanding of actionable genomic alterations with 50–60% of tumors harboring *BRAF* mutations (*1*), and the development of effective therapies, including targeted BRAF/MEK inhibition (*2*) and immunotherapy (immune checkpoint blockade) (*3*). Despite these advances, durable benefits remain limited to a subset of patients (*2–4*). Among individuals with *BRAF*-mutant metastatic melanoma treated with combined BRAF/MEK inhibition (BRAFi/MEKi), approximately 20% remain progression-free survival (PFS) at 5 years (*4*), while long-term overall survival following PD-1-based immunotherapy is observed in 40–60% of *BRAF*-mutant melanoma patients (*5*).

Recent phase III clinical trials, including DREAMseq (*6*) and SECOMBIT (*7*), have addressed the optimal sequencing of BRAF-targeted therapy and immunotherapy. At a cohort level, these studies demonstrated superior survival when combination immunotherapy (nivolumab plus ipilimumab) is administered prior to BRAF-targeted therapy (dabrafenib plus trametinib), compared with the reverse sequence. However, it remains unclear whether outcomes can be improved by precision approaches to identify and potentially target the patients who would have a durable response to targeted therapies.

Although the initial response rates to BRAFi/MEKi are high (60–70%) in *BRAF*-mutant metastatic melanoma, durable responses remain uncommon, and the baseline predictors of long-term benefit are poorly defined. Resistance-associated genomic alterations, including mitogen-activated protein kinase (MAPK) pathway reactivation through *MEK1/2* or *ERK* mutations and *CDKN2A* loss, have been well characterized in post-treatment tumors and experimental models (*8–10*), but their presence in pretreatment samples does not appear to predict therapeutic efficacy (*8*). Exploratory analyses of baseline tumors from *BRAF*-mutant patients treated with BRAFi/MEKi have identified candidate alterations associated with complete response (e.g., *NF1*) and progression (e.g., *MITF* and *TP5*3) (*11*). However, the low frequency of these alterations in pretreatment samples does not explain the majority of rapid progression cases. In addition, the absence of long-term clinical follow-up in this study precludes the evaluation of baseline features associated with durable benefit from BRAF-targeted therapy.

Beyond tumor-intrinsic features, accumulating evidence suggests that the tumor immune microenvironment may influence response to BRAF-targeted therapy. Oncogenic BRAF signaling has been shown to promote immune suppression in preclinical models, while BRAFi and MEKi can transiently enhance tumor immunogenicity and immune infiltration (*12–14*).

Consistent with this, transcriptomic and histologic studies of pretreatment tumors from *BRAF*-mutant metastatic melanoma have reported associations between immune-related gene expression (*11*), CD8^+^ T cell density (*15*), and improved clinical outcomes following BRAFi/MEKi targeted therapy. However, the baseline immune features, and their spatial organization, that distinguish patients with durable benefit from those with early resistance remain incompletely understood.

In this study, we systematically investigated baseline tumor-intrinsic and microenvironmental features in a subset of patients enrolled in a large phase III clinical trial of BRAFi/MEKi with long-term follow-up (COMBI-d) (*4*). By stratifying patients into distinct response groups, we identified tumor genomic heterogeneity and tumor burden as key features to distinguish these groups in response to BRAF-targeted therapy. Building on these findings, we developed a parsimonious predictive model to identify patients with durable response or resistance with high precision and specificity. Notably, patients predicted to have durable responses to targeted therapy did not exhibit durable responses in anti–PD-1-treated cohorts(*16–18*), suggesting that distinct patient subgroups may preferentially derive long-term benefit from BRAF-targeted therapy versus immune therapy. In parallel, we characterized three distinct tumor microenvironments based on CD8^+^ T cell spatial architecture, with high density intratumoral (but not peritumoral) CD8^+^ T cell infiltration associated with durable response to BRAF-targeted therapy.

## RESULTS

### Genomic and clinical characterization of the COMBI-d cohort with survival based stratification

Whole-exome sequencing (WES) was performed on pre-treatment tumor tissue and matched blood from a total of 167 patients in the COMBI-d trial (*4*). After quality control (Materials and Methods), WES data from 157 patients were included in this study (∼37% of the COMBI-d trial; **fig. S1A**). Demographic and baseline characteristics of the WES cohort were similar to those of the full intention-to-treat (ITT) trial population (**fig. S2A** – Therapy, **table S1**) (*4*). *BRAF (*p.V600E) mutations, initially detected by the qPCR-based assay (FDA-approved bioMérieux BRAF THxID assay kit) (*19*) at the time of screening, were confirmed in almost all baseline samples with available WES data (155/157) (**fig. S1A**, **fig. S2A** – *BRAF* Mutation Subtype). The two patients without *BRAF* mutations detected were excluded from further analysis. Best response (BR) to BRAF-targeted therapy using RECIST (v.1.1) criteria (Methods) included 11% progressive disease (PD), 22% stable disease (SD), 43% partial response (PR), and 21% complete response (CR) (**fig. S2A** - Best Response).

Evaluation of PFS revealed a distinct subgroup of patients with durable benefit (**Fig. 1A**), which we categorized as “durable responders” (DR; PFS ≥ 24 months; n = 39). We further characterized “rapid progressors” (RP; n = 30) based on PFS ≤ 6 months and progressed (**Fig. 1A**). The rest of the patients were defined as “intermediate outcomes” (INT; n = 86). For the RP patient group, we excluded patients treated with BRAFi alone, as it was unclear whether they would have remained RP if treated with BRAFi/MEKi combination therapy. For the DR population, patients treated with BRAFi monotherapy were included, under the assumption that DR to BRAFi would also be DR to BRAFi/MEKi combination therapy.

**Figure 1.**
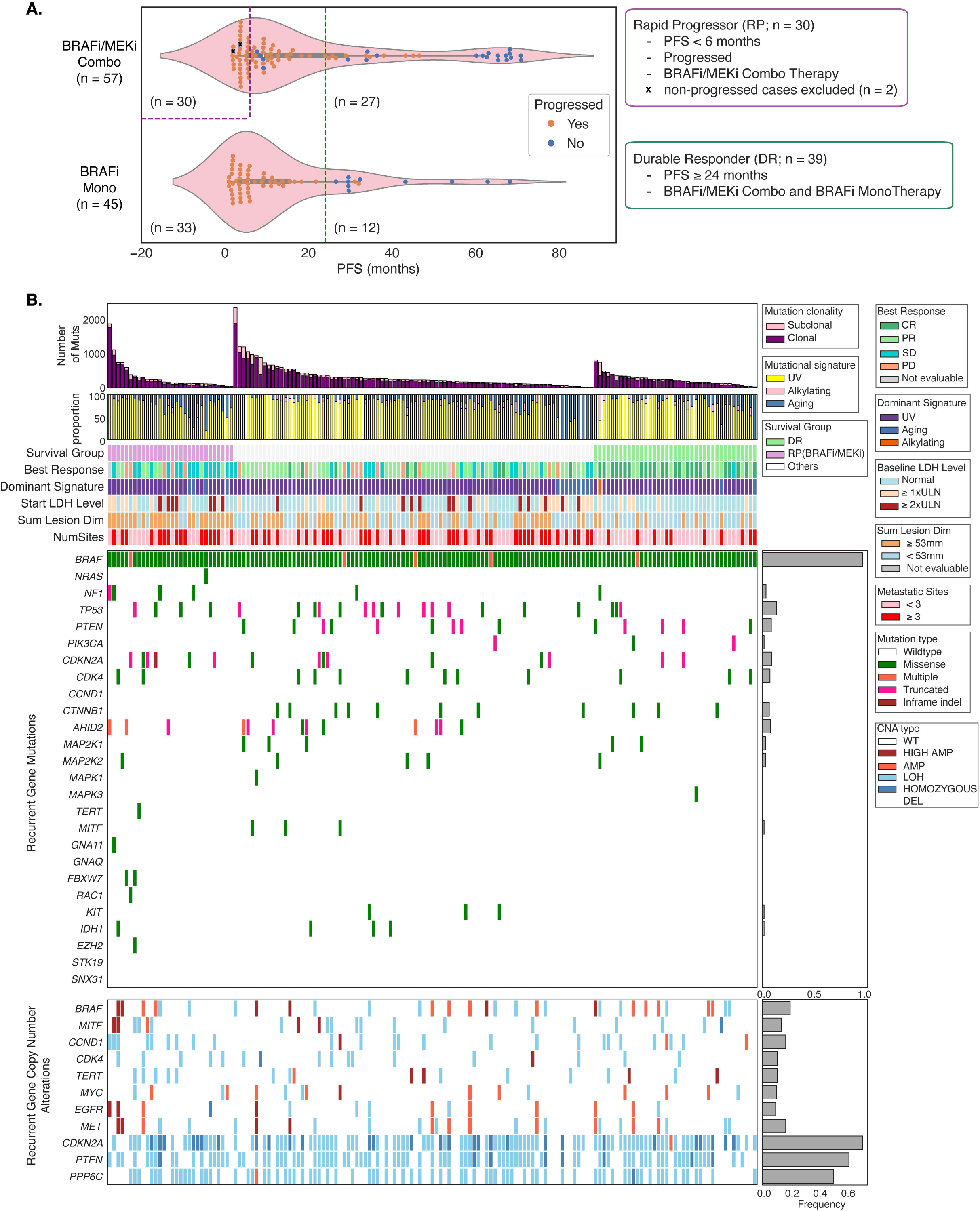
Overview of clinical outcomes and genomic features in the COMBI-d cohort. (**A**) Violin plots showing PFS of metastatic melanoma patients treated with BRAFi/MEKi combination therapy and BRAFi monotherapy. Orange dots represent patients who progressed. The purple dashed line indicates the threshold defining RP (PFS < 6 months with progression), while the green dashed line indicates the threshold defining DR (PFS ≥ 24 months). Crossed blue dots indicate patients who did not progress and were excluded from the RP group despite having PFS < 6 months (n = 2). (**B**) CoMut plot showing the association between clinical and genomic characteristics. Each column represents a tumor. The columns were ordered by DR, INT, and RP within each response subgroup by decreasing nonsynonymous mutational load (top row). Nonsynonymous mutational burden is further subdivided into clonal (purple) and subclonal (pink) mutational loads. Mutational signatures refer to the inferred relative contribution of UV induced mutations (yellow), alkylating DNA damaging–associated process (pink) and aging mutational signatures (blue). Best response was defined by RECIST criteria response: CR (green), PR (light green), PD (orange), SD (blue). Durable response was defined as PFS ≥ 24 months (light green). The start level of lactate dehydrogenase (LDH) was classified as normal (grey), ≥ 1× ULN (light brown), and ≥ 2× ULN (brown). Sum of lesion dimensions refers to the inferred total size of metastatic lesions, grouped by equal or above (orange) and below (blue) the median value (53 mm); two patients could not be evaluated (gray). The metastatic sites represent the total number of metastatic lesions, categorized as equal or above (red) and below (pink) three. Mutations in *BRAF*, *NRAS*, *NF1*, Cell cycle–associated genes (e.g., *TP53*, *CDKN2A*, *CDK4*, *CCND1*), mTOR pathway–associated genes (e.g., *PIK3CA*, *PTEN*), MAPK pathway–associated genes (*MAP2K1*, *MAP2K2*, *MAPK1*, *MAPK3*) and other known cancer-related genes were shown in each tumor. Copy number alterations in those recurrent cancer genes are shown in the bottom panel. Amplifications (red), high-level amplifications (dark red), loss of heterozygosity (light blue), and homozygous deletions (dark blue) are indicated. The right bar plot summarizes the frequency of copy number alterations across the cohort.

To characterize the most frequently mutated genes in our WES data, we investigated mutations and copy number alterations in well-known tumor suppressors and other previously described genetic alterations in TCGA and other melanoma sequencing datasets (**Fig. 1B**, **table S2**) (*1, 20*). Besides *BRAF*, *TP53* was the most frequently altered gene (14%), following by *CDKN2A* (10%) and *PTEN* (9%) in the COMBI-d cohort (**Fig. 1B –** Recurrent Gene Mutations). Most of the patients had deletions in tumor suppressor genes such as *CDKN2A* and *PTEN* (**Fig. 1B –** Recurrent Gene Copy Number Alterations). As expected for a cutaneous melanoma cohort, most patients had a dominant UV signature. Alkylating and aging signatures dominated over the UV signature in 1 and 15 tumors, respectively (**Fig. 1B –** Dominant Signatures). The one patient with dominant alkylating signature (*21*) had prior treatment with chemotherapy. However, none of the recurrent gene mutations, copy number alterations, or mutational signatures were enriched within either DR or RP patient groups.

### Distinct genomic and clinical feature axes characterized durable response in the COMBI-d cohort

We next directly compared the overall genomic and clinical features between DR, INT, and RP. Consistent with previous findings (*11*), there was no difference in tumor mutational burden (**Fig. 2A** – Nonsyn Mutation Load). In contrast, RP exhibited significantly higher tumor genomic heterogeneity than DR (MWW, *P* = 0.036) (**Fig. 2A** – Heterogeneity). Although melanoma evolution studies have suggested that late-stage metastatic melanoma is dominated by aneuploidy and whole genome doubling (*22*), and a recent study showed that ploidy is associated with intrinsic resistance to immunotherapy in metastatic melanoma (*23*), we found no significant difference in ploidy between RP and DR (MWW, *P* = 0.73) (**Fig. 2A** – Ploidy) or in the proportion of the tumor genome with aneuploidy (MWW, *P* = 0.88) (**Fig. 2A** – Copy Number Alteration). A trend toward lower tumor purity was observed in DR (MWW, *P* = 0.061) (**Fig. 2A** – Purity), suggesting a more immune-infiltrated tumor microenvironment in durable responders to BRAF-targeted therapy.

**Figure 2.**
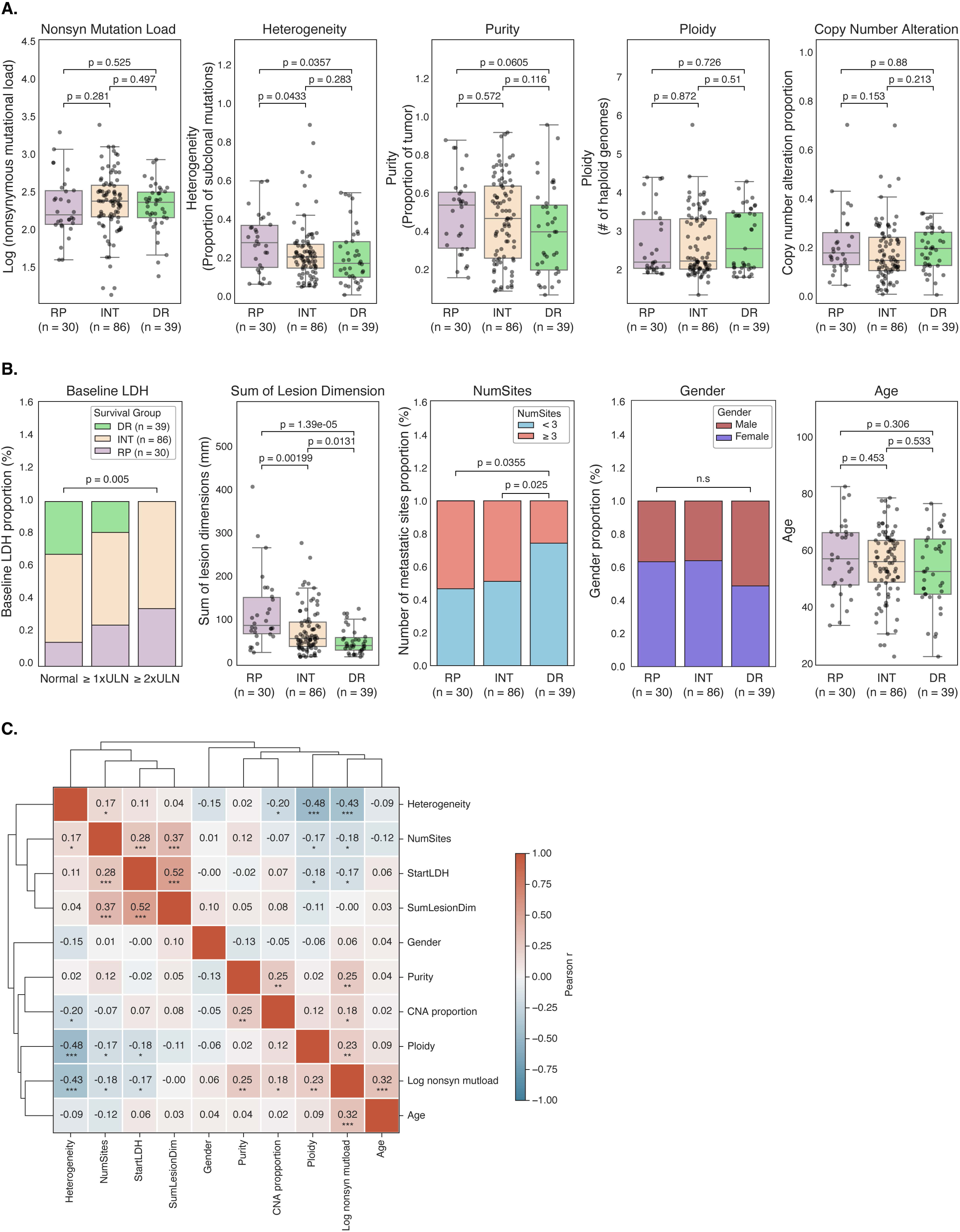
Distinct genomic and clinical feature axes characterize durable response in the COMBI-d cohort. **(A)** Boxplots showing genomic features comparison between RP (n = 30), INT (n = 86), and DR (n = 39) groups. Shown are log-transformed nonsynonymous mutational load, intratumoral heterogeneity, purity, ploidy and copy number alteration proportion. Group comparisons were performed using two-sided Mann-Whitney-Wilcoxon tests with Bonferroni correction; corresponding P values are indicated in the figure. **(B)** Boxplots showing clinical comparison across RP (n = 30), INT (n = 86) and DR (n = 39) groups. Shown are sum of lesion dimensions, number of metastatic sites, baseline LDH level, sex, and age at treatment initiation. Group comparisons were performed using two-sided Mann–Whitney–Wilcoxon tests for continuous variables and chi-squared tests for categorical variables; corresponding P values are indicated in the figure. **(C)** Heatmap showing pairwise Pearson correlation coefficients among genomic and clinical features. Features are ordered by hierarchical clustering based on correlation distance. Pearson correlation coefficients are shown within each cell, with asterisk annotation indicating nominal significance levels as shown. Color scale represents the strength and direction of correlation. Asterisks denote nominal significance (*: P < 0.05; **: P < 0.01; ***: P < 0.001).

Clinically, tumors from RP were associated with poor prognostic features related to tumor burden, including higher baseline LDH levels (χ², *P* = 0.005, **Fig. 2B** – Baseline LDH), greater sum of lesion dimensions (MWW, *P* < 0.0001) (**Fig. 2B** – Sum of Lesion Dimension), and a higher number of metastatic sites (χ², *P* = 0.035) (**Fig. 2B** – NumSites). In contrast, no significant differences were observed between survival groups with respect to sex (χ², *P* = 0.33) (**Fig. 2B** – Sex) and age (MWW, *P* = 0.31) (**Fig. 2B** – Age).

To understand the relationships among features associated with response, we evaluated correlations among features and clustered them based on their similarities. Clinical features associated with tumor burden, including sum of lesion dimensions, number of metastatic sites, and baseline LDH, were strongly correlated and clustered together (**Fig. 2C**). In contrast, tumor genomic heterogeneity was largely uncorrelated with tumor burden (correlation coefficients of 0.04–0.17) and was weakly negatively correlated with mutation burden, ploidy, and CNA proportion (correlation coefficients of –0.43, –0.48, and –0.2, respectively).

### Gene-level correlates of durable response and rapid progression

Given the observed genomic heterogeneity across tumors, we performed an unbiased analysis to identify functional single-gene alterations associated with DR or RP to BRAF-targeted therapy (*16, 24*). Gain-of-function (GOF) gene alterations were defined as gene amplification or the presence of GOF hotspot mutations annotated by OncoKB (Methods) (*25*), while loss-of-function (LOF) gene alterations were defined as biallelic gene-level inactivation events, including homozygous deletions or a combination of functionally inactivating mutations (Methods).

Among recurrently amplified genomic regions, amplification at chromosome 11q13.1 including *LTBP3* (latent transforming growth factor beta-binding protein 3) was most strongly associated with DR (11/38 DR versus 1/30 RP; Fisher’s exact test, *P* = 0.0085, adjusted odds ratio = 6.43) (**fig. S3A**). In total, 17 patients in the cohort harbored *LTBP3* amplification (**fig. S3B, table S3**), and had longer PFS compared with patients without *LTBP3* amplification (**fig. S3E**). While there were no significant differences in baseline clinical features between patients with and without *LTBP3* amplification (**fig. S3D**), those with *LTBP3* amplification had lower heterogeneity, increased ploidy, and a trend toward lower tumor burden (**fig. S3C**). We then evaluated the presence of *LTBP3* amplifications in other metastatic melanoma cohorts (*16, 23, 26–31*) with exome- or genome-wide copy number alterations, and observed this in 2–6% of metastatic melanoma populations (**fig. S3F**).

The strongest correlate of RP within genomic amplifications was chromosome 3p14.1 including *MITF,* which was previously nominated as a mechanism of resistance in preclinical and clinical studies (*8, 11, 32*), although this did not reach statistical significance in our size-limited cohort (0/38 DR versus 3/30 RP; Fisher’s exact test, *P* = 0.08, adjusted odds ratio = 0.18) (**fig. S3A, table S3**). Two additional patients in the COMBI-d cohort harbored *MITF* amplification; although neither met criteria for RP, both experienced disease progression within one year of therapy (**fig. S3B**).

To further evaluate whether *MITF* amplification could serve as a biomarker of RP, we analyzed an independent cohort of 22 patients with *BRAF*-mutant metastatic melanoma treated with BRAF-targeted therapy, whose tumors harbored *MITF* amplification as identified by a targeted gene panel assay (*34*). In this cohort, while ∼36% (8/22) were RP, we also observed ∼27% (6/22) as DR (**fig. S4A**). Among the subset of patients with high amplification of *MITF* (n = 3), two were RP and one was DR (**fig. S4A**). Finally, we evaluated patients with versus without *MITF* amplification in the combined cohort and found no significant difference in PFS between these groups (**fig. S4B**).

In the LOF analysis, we identified regions of chromosome 8, 19 and 1 associated with RP (**Chr 8, 19**) and DR (**Chr 1**) (**fig. S4C, table S4**). However, the functional significance of the genes from those regions in resistance to BRAF-targeted therapy is unclear.

### A parsimonious predictive model identified tumor heterogeneity and tumor burden as treatment-specific determinants of response to BRAF-targeted therapy

Given the observed genomic and clinical differences between DR and RP (**Fig. 2A, B**), we hypothesized that integrating genomic heterogeneity with tumor burden-associated features could stratify PFS. Indeed, stratifying patients into four groups using median splits of genomic heterogeneity and sum of lesion dimensions significantly separated PFS, with the most favorable outcomes observed in patients with low heterogeneity and low sum of lesion dimensions, whereas patients with high heterogeneity and high sum of lesion dimensions were associated with the worst PFS (global log-rank *P* = 0.0013, **Fig. 3A**). We observed similar results when using baseline LDH and number of metastatic sites as metrics of tumor burden with genomic heterogeneity (**fig. S5A, B**).

**Figure 3.**
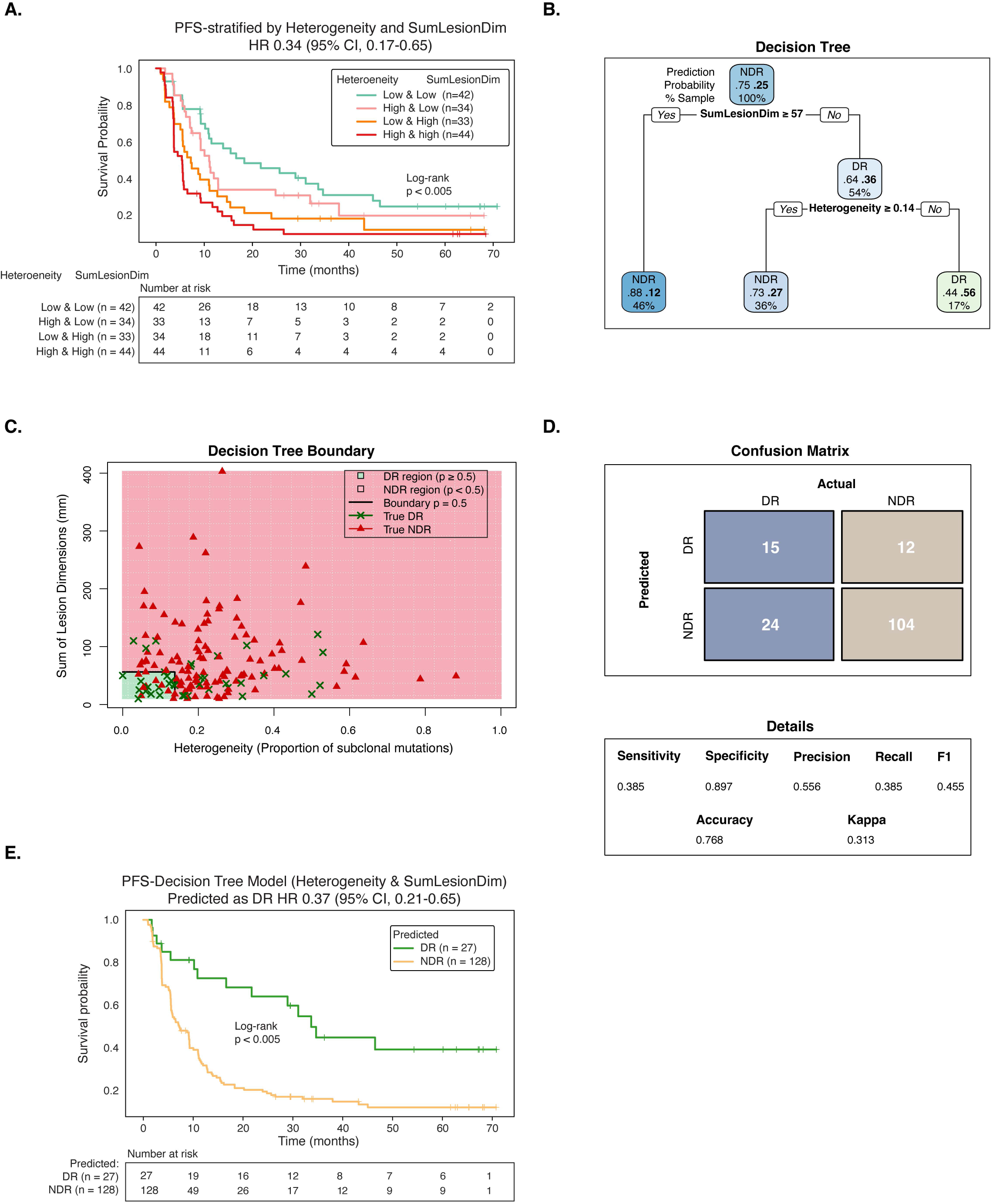
Decision tree models based on tumor heterogeneity and sum of lesion dimensions to predict RP in the COMBI-d cohort. **(A)** Kaplan–Meier PFS curves stratifying patients by median dichotomization of sum of lesion dimensions and heterogeneity into four groups. P values were calculated using the log-rank test. **(B)** Structure of the decision tree model used to classify DR versus NDR based on tumor genomic heterogeneity and sum of lesion dimensions. For each node, the predicted class is shown at the top, the estimated probabilities of NDR (left) and DR (right) are shown in the middle, and the percentage of patients assigned to the node is shown at the bottom. **(C)** Decision boundaries defined by thresholds on heterogeneity and sum of lesion dimensions. Background shading indicates the model-predicted probability of DR, where p = P(DR); p ≥ 0.5 is classified as DR and p < 0.5 as NDR. Individual patients are overlaid, with green crosses denoting true DR and red triangles denoting true NDR. **(D)** Confusion matrix summarizing model performance in the cohort, showing predicted versus observed response categories. **(E)** Kaplan–Meier PFS curves stratified by decision tree–predicted DR and NDR groups in the COMBI-d cohort. P values were calculated using the log-rank test.

Next, we tested whether these features could be leveraged to develop predictive models to identify patient subgroups enriched in DR. Without any selection, 25% of the entire cohort were DR. Utilizing clinical metrics of tumor burden (baseline LDH, sum of lesion dimensions, number of metastatic lesions) in a single feature decision tree (Methods), we identified patient subgroups with 38–44% DR (**fig. S5C–F**). Integrating genomic heterogeneity with metrics of tumor burden further improved model performance, enabling identification of patients further enriched for DR (52–56%) (**Fig. 3B–D**, **Fig. 4A–C**, **fig. S6A–C,** and **fig. S6E**). The predicted patients had significantly improved PFS compared with the overall cohort (**Fig. 3E**, **Fig. 4D**, and **fig. S6D**).

**Figure 4.**
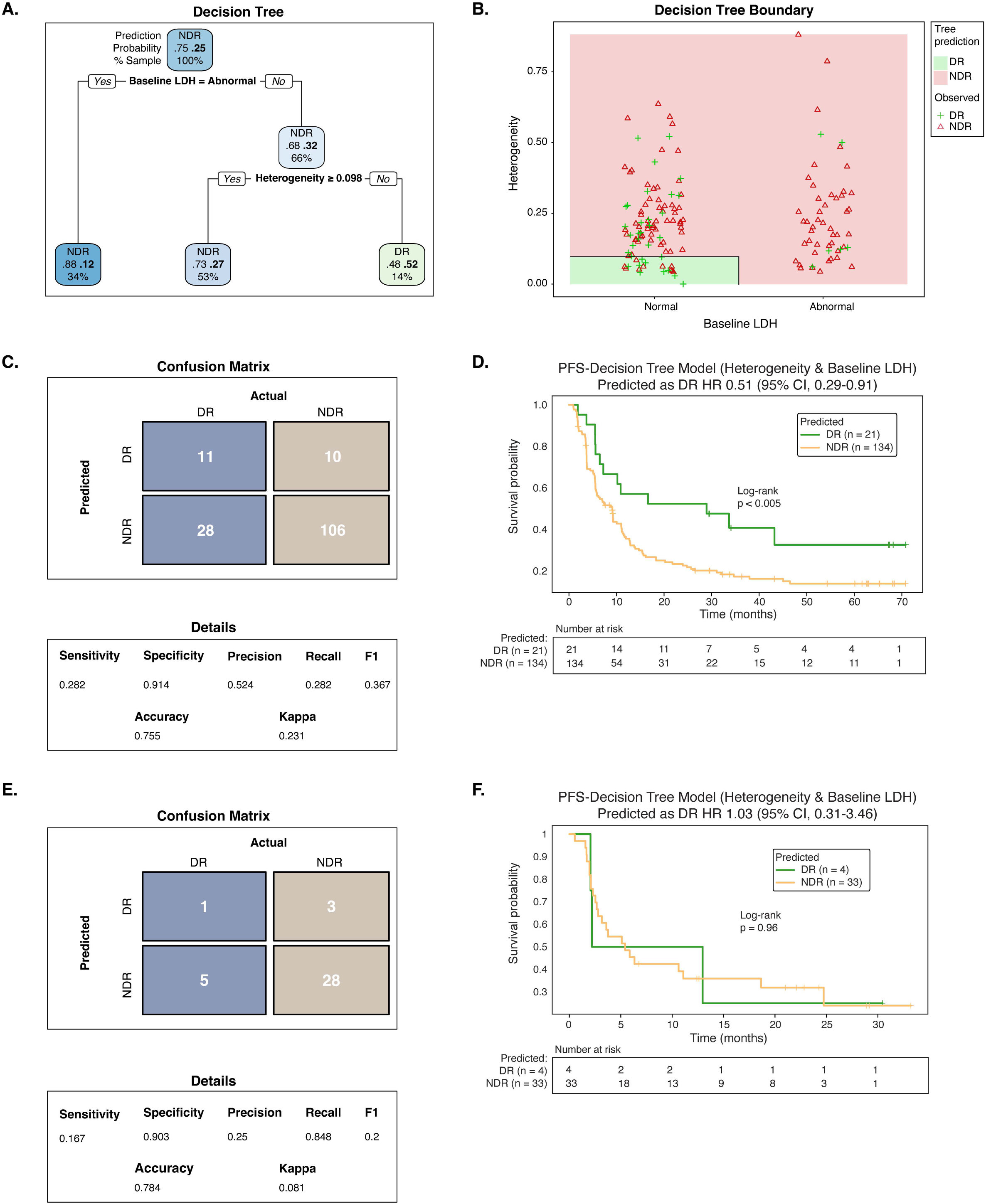
Decision tree models based on tumor heterogeneity and baseline LDH to predict DR and NDR in the COMBI-d cohort. **(A)** Structure of the decision tree model used to classify DR versus NDR based on tumor genomic heterogeneity and baseline LDH. For each node, the predicted class is shown at the top, the estimated probabilities of NDR (left) and DR (right) are shown in the middle, and the percentage of patients assigned to the node is shown at the bottom. **(B)** Decision boundaries are defined by thresholds on tumor genomic heterogeneity and baseline LDH status. Background shading indicates the model-predicted probability of DR, where p = P(DR); p ≥ 0.5 is classified as DR and p < 0.5 as NDR. Individual patients are overlaid, with green crosses denoting true DR and red triangles denoting true NDR. **(C)** Confusion matrix summarizing model performance in the cohort, showing predicted versus observed response categories. **(D)** Kaplan–Meier PFS curves stratified by decision tree–predicted DR and NDR groups in the COMBI-d cohort. P values were calculated using the log-rank test. **(E)** Confusion matrix summarizing the application of the decision tree model to the anti–PD-1 cohort after excluding patients with PFS ≤ 24 months without progression, showing predicted versus observed response groups. **(F)** Kaplan-Meier PFS curves stratified by decision tree–predicted DR and NDR groups in the anti–PD-1 cohort. P values were calculated using the log-rank test.

To determine whether these features were prognostic or treatment-specific, we applied the model trained on tumor heterogeneity and baseline LDH from the COMBI-d cohort (**Fig. 4A**–**D**) to a harmonized anti–PD-1-treated cohort of patients with *BRAF*-mutant metastatic melanoma (n = 38) who had not received prior BRAF-targeted therapy, assembled from previous studies and clinical trials (*16–18*). No significant differences in heterogeneity or the proportion of abnormal baseline LDH were observed between cohorts (**fig. S7A, B**). The model performed poorly in this setting: of four patients predicted to be DR, only one was a true DR (**Fig. 4E**), and PFS did not differ between predicted DR and non-DR (**Fig. 4F**) in this limited cohort. Consistently, no difference in PFS was observed when we stratified patients from the anti–PD-1-treated cohort into four groups based on the median values of tumor genomic heterogeneity and baseline LDH (**fig. S7C**). These findings suggest that the predictive features identified for BRAF-targeted therapy are treatment-specific and that DR to BRAF-targeted therapy and anti–PD-1 immunotherapy are characterized by distinct baseline features.

### Distinct spatial CD8^+^ T cell immune phenotypes characterized response to BRAF-targeted therapy

Growing evidence supports a role for tumor immunity in the therapeutic efficacy of BRAF-targeted therapy in metastatic melanoma (*11, 15*). Genomic analysis of the COMBI-d cohort revealed a trend toward lower tumor purity in DR compared with RP (**Fig. 2A –** Purity), suggesting increased non-tumor cell content. To assess whether features of the tumor microenvironment are associated with response to BRAF-targeted therapy, we analyzed CD8 immunohistochemistry (IHC) data from available pretreatment tumor samples in the COMBI-d cohort (n = 47). Considering the tumor as a whole, we observed a trend toward lower CD8^+^ T cell density in the RP tumors compared with INT tumors, but similar levels when compared with DR tumors (**fig. S9A**).

To further assess whether the specific spatial distribution of CD8^+^ T cells is associated with response to BRAF-targeted therapy, we performed a spatial band analysis quantifying CD8^+^ T cell density at the invasive margin (Methods). CD8^+^ T cell density was measured in consecutive 30-μm bands extending up to 150 μm into the tumor (intratumoral) and 150 μm into the adjacent stroma from the tumor boundary (**Fig. 5A**, **fig. S8A, B**). Based on the spatial localization of CD8^+^ T cells, tumors were classified into three immune phenotypes: inflamed (high CD8^+^ T cell density in both intratumoral and peritumoral regions; n = 20), desert (low CD8^+^ T cell density in both intratumoral and peritumoral regions; n = 14), and excluded (high peritumoral but low intratumoral CD8^+^ T cell density; n = 13) (**Fig. 5A**, **table S5**). Hierarchical clustering independently recapitulated these immune phenotypes (**fig. S9C**), which were not associated with biopsy sites (χ², P = 0.25) (**fig. S9B**).

**Figure 5.**
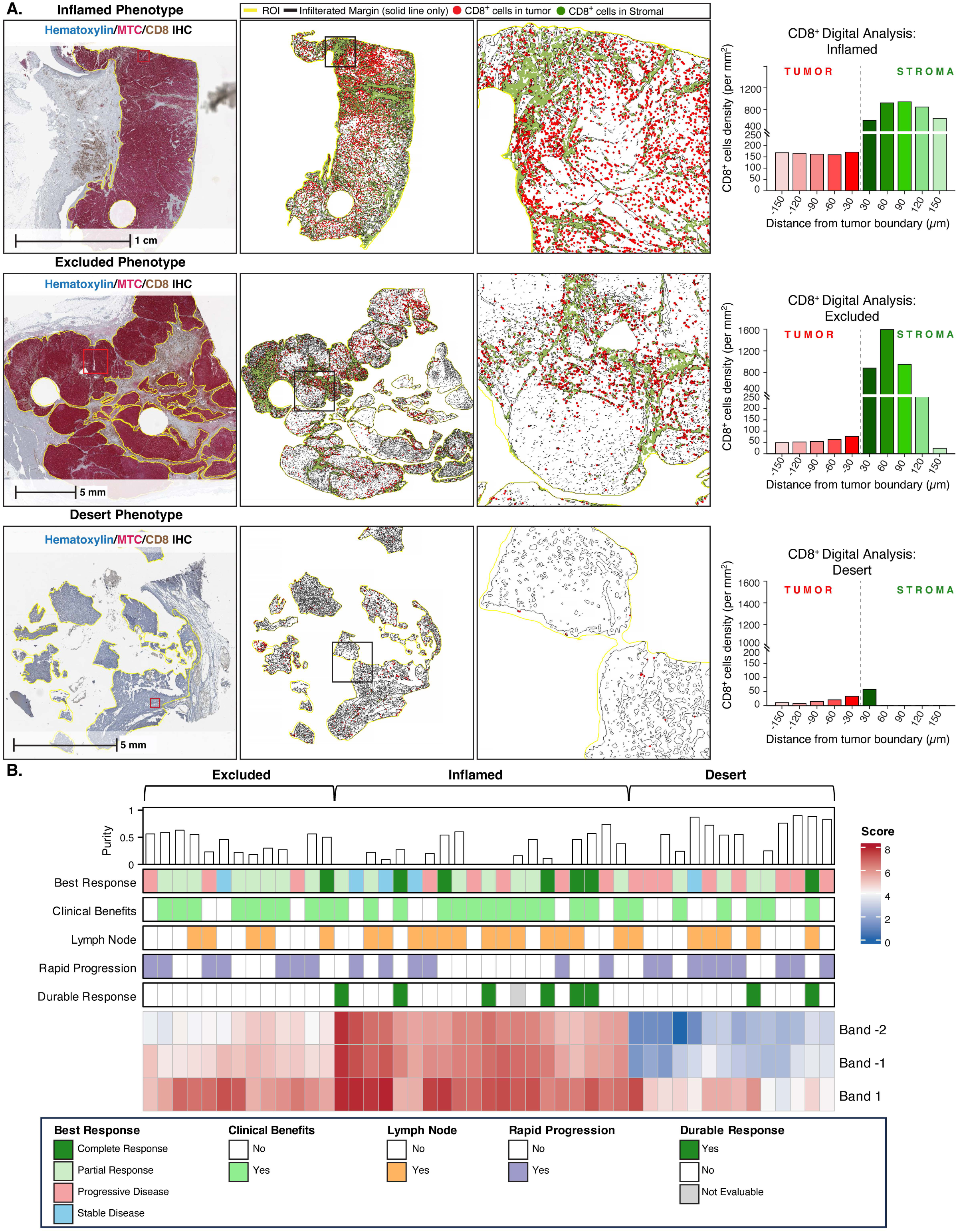
Spatial distribution of CD8^+^ T cells within the COMBI-d cohort. **(A)** Representative images illustrating inflamed, excluded, and desert tumor-immune phenotypes, defined by the spatial distribution of CD8⁺ T cells relative to intratumoral and stromal compartments at the invasive margin. *Column 1*: Dualplex chromogenic IHC of whole-slide tumor sections stained with hematoxylin (blue), melanoma triple cocktail (MTC; red), and CD8 (brown). Tumor regions for the entire tissue lesion were annotated by a pathologist based on histomorphological assessment, using MTC-positive melanoma tumor cells as a reference. The yellow outline indicates the tumor boundary (invasive margin), which was used as the infiltration margin reference for spatial proximity analyses. Red boxed regions correspond to areas shown at higher magnification in Fig. S8A. Scale bars, 1 and 5 mm. *Column 2*: Spatial scatter plots depicting the distribution of CD8⁺ T cells across the tissue section. CD8⁺ T cells are colored red within melanoma tumor lesions and green within stromal regions. Yellow lines denote tumor regions (regions of interest, ROIs), and black lines indicate the infiltrating margin. Black boxes highlight regions shown at higher magnification in *Column 3*. Scale bars, 1 and 5 mm. *Column 3*: High-magnification views illustrating the spatial localization of CD8⁺ T cells within intratumoral and stromal compartments. *Column 4*: Quantification of CD8⁺ T cell density (cells/mm²) across spatial bands relative to the tumor boundary for the corresponding samples showing in Columns 1–3. Negative distances represent intratumoral regions, whereas positive distances represent stromal regions. Bars are colored to reflect spatial location and CD8^+^ T cell density, with red gradients indicating intratumoral regions and green gradients indicating stromal regions. Data are shown for individual patients representing each phenotype: inflamed (G053), excluded (G023), and desert (G073). **(B)** CoMut plot summarizing CD8⁺ T cell infiltration patterns alongside clinical and treatment-related features. Each column represents an individual tumor. Best response is defined according to RECIST criteria (CR, PR, SD, and PD). Lymph node denotes tumors sampled from lymph nodes (orange) or non–lymph node sites (white). Rapid progression is defined as PFS < 6 months with progression, and durable response as PFS ≥ 24 months. Band rows indicate CD8^+^ T cell infiltration patterns. Band 1 denotes the peritumoral region extending outward from the tumor boundary, while bands −1 and −2 denote intratumoral regions extending inward; each band represents a 30-µm interval.

Notably, there were no DR among patients with an excluded immune phenotype (**Fig. 5B**). Consistent with this observation, RP exhibited significantly lower intratumoral CD8^+^ T cell density compared with non-RP (MWW, *P* = 0.027) (**Fig. 6A**), while DR showed a trend toward higher intratumoral, but not peritumoral, CD8^+^ T cell density compared with non-DR (**fig. S9D, E**). Moreover, patients with inflamed tumors had significantly prolonged PFS compared with all other immune phenotypes (**Fig. 6B**). Although prior studies reported enrichment of CD8⁺ T effector, cytolytic T cell, antigen presentation, and natural killer cell gene signatures in complete responders to targeted therapy (*11*), these findings suggest that intratumoral CD8⁺ T cell localization, rather than overall abundance, is a key determinant of response to BRAF-targeted therapy. Notably, intratumoral CD8^+^ T cell infiltration was independent of genomic heterogeneity and tumor burden associated features (**fig. S9F**).

**Figure 6.**
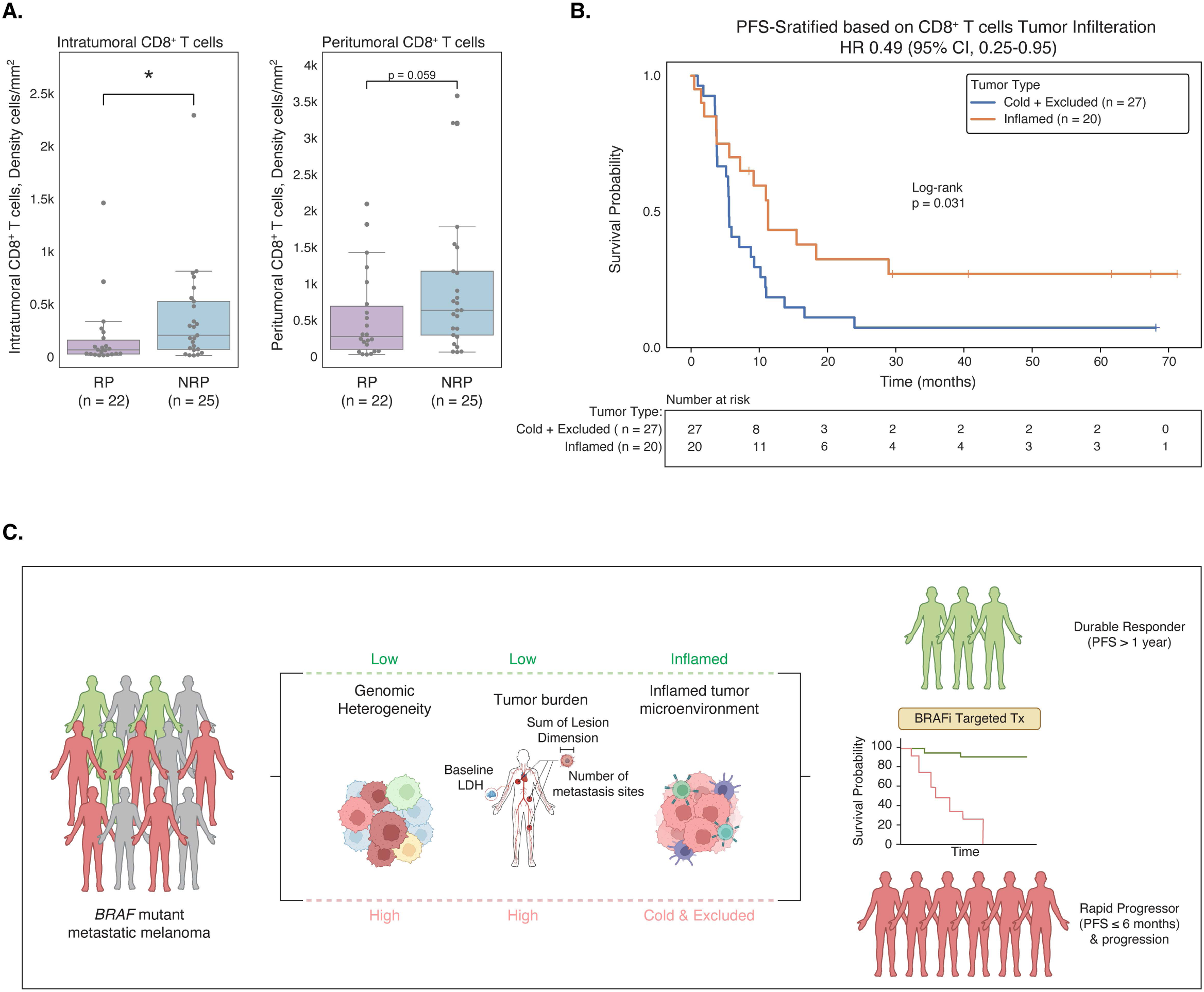
Intratumoral CD8^+^ T cells are associated with response to BRAF-targeted therapy. **(A)** Comparison of CD8⁺ T cell density between RP and non-rapid progressors (NRP; PFS ≥ 6 months). Intratumoral CD8⁺ T cell density was calculated as the mean density across bands −5 to −1, and peritumoral CD8⁺ T cell density as the mean density across bands 1 to 5. Group comparisons were performed using a two-sided Mann–Whitney–Wilcoxon test with Bonferroni correction; corresponding P values are shown. **(B)** Kaplan–Meier PFS curves stratified by tumor immune phenotype (inflamed versus excluded/desert). P values were calculated using the log-rank test. **(C)** Schematic summary of genomic, clinical, and immune features that stratify DR in BRAF-targeted therapy.

Together, our data support the hypothesis that durable responders to targeted therapy harbor distinct tumor-intrinsic and extrinsic features (**Fig. 6C**), towards the development of clinically actionable biomarkers for individual patients.

## DISCUSSION

In this study, we investigated pre-treatment predictors of durable response and resistance to BRAF-targeted therapy in a large, clinically annotated cohort of patients with *BRAF*-mutant metastatic melanoma with long-term follow-up, and identified multiple genomic, clinical, and spatial features distinguishing durable responders and rapid progressors. We further found that these features provide non-overlapping information and can be integrated to identify groups of patients enriched for durable responders with substantially improved PFS.

Intratumoral genomic heterogeneity has been associated with poor prognosis across multiple tumor types and therapies (*35*). High heterogeneity reflects a highly mutagenic disease state with increased subclonal diversity, thereby increasing the likelihood of preexisting or rapidly emerging resistant clones. Consistent with this, we found that DR had significantly lower genomic heterogeneity compared with RP, suggesting that greater subclonal diversity predicts the rise of resistant clones. Conversely, tumor mutational burden does not predict survival, DR, or RP.

In individual gene mutation analyses, baseline amplification of *LTBP3* was significantly associated with DR to BRAF-targeted therapy, and tumors harboring *LTBP3* amplification exhibited distinct genomic features, particularly with low intratumoral heterogeneity compared to those without amplification. Although the role of *LTBP3* amplification in metastatic melanoma is not well understood, LTBP3 is a critical regulator of transforming growth factor-β (TGF-β) (*36*). TGF-β signaling has a complex role in BRAF-targeted therapy. While it can promote resistance to BRAF-targeted therapy in melanoma by driving pro-invasive, pro-survival pathways, a recent study showed that high TGF-β activation, in combination with high dose MAPK inhibition, can paradoxically induce apoptosis by shifting TGF-β’s effect from pro-tumorigenic to pro-apoptotic (*37*). We hypothesize that *LTBP3* amplification may modulate TGF-β signaling at an early stage, thereby reducing tumor invasiveness and contributing to durable response to BRAF-targeted therapy. Further work is necessary to functionally validate the biological mechanisms of *LTBP3* amplification and to independently assess its potential as a biomarker for durable responders to BRAF-targeted therapy.

While the association between *MITF* amplification and resistance to BRAF-targeted therapy remains controversial, preclinical studies have shown that upregulation of MITF, potentially regulated by *PAX3* and its downstream targets (e.g., *MLANA*, *TYRP-1*, *PMEL*), is observed in highly proliferative melanoma cells and has been connected to long-term resistance to BRAF-targeted therapy (*38*). Consistent with this model, a prior correlative analysis from a clinical trial reported an enrichment of *MITF* amplification among patients with progressive disease (PD), with 15% of PD cases harboring *MITF* amplification (11/70) (*11*).

In our cohort, *MITF* amplification was also more frequently observed among RP compared with DR tumors. Although this association is not statistically significant, likely due to the small number of patients with *MITF* amplification in our cohort (10% of RP; 3/30 RP, n = 5 patients with *MITF* amplification in total), the proportion observed in our study is comparable to the prior report (*11*).

However, comparison of PFS between patients with and without MITF amplification in the COMBI-d and DFCI validation cohorts revealed no significant difference. Given the limited sample size, further validation in larger, independent cohorts will be required to determine whether and in what context *MITF* amplification is a biomarker of rapid progression in BRAF-targeted therapy.

Towards predicting durable responders, simple models integrating clinical metrics of tumor burden and genomic heterogeneity were able to identify subgroups of patients enriched for durable responders (52–56% versus 25% for the unselected cohort). Baseline metastatic growth rate, including elevated LDH, metastatic sites, and tumor growth rate, has previously been implicated as a prognostic marker in patients with advanced *BRAF^V600^*-mutant melanoma receiving BRAF targeted therapy (*39*) and immunotherapy (*40*). However, the poor performance of this model in predicting durable response in an independent cohort of *BRAF*-mutant melanoma patients treated with anti–PD-1 immunotherapy suggests that the set of durable responders to targeted therapy may be distinct from patients with durable response to immune checkpoint blockade.

It is intriguing to consider whether genomic heterogeneity in a very low tumor burden setting (e.g. in resected Stage III disease) would predict durable response and cure with adjuvant targeted therapy. Unlike unresectable Stage III and IV disease, there are currently no randomized studies guiding the optimal choice of adjuvant therapy in resected advanced melanomas, and our data suggest that tumors with low genomic heterogeneity may derive disproportionate benefit from targeted therapy in this setting.

Recent evidence indicates that the tumor microenvironment plays an important role in modulating response and resistance in BRAF-targeted therapy. Oncogenic BRAF has been shown to promote immune evasion in melanoma through multiple mechanisms, including the secretion of immunosuppressive cytokines (*41*), modulation of major histocompatibility complex (MHC) expression in tumor cells (*42*), and downregulation of CCL2 expression and decreased tumor CCL2 protein release (*43*). Conversely, inhibition of oncogenic BRAF through MAPK pathway blockade has been associated with increased expression of melanoma differentiation antigens, suggesting that BRAF-targeted therapy may enhance tumor immunogenicity (*44*).

Consistent with this concept, pooled analyses of multiple BRAFi/MEKi clinical trials have demonstrated that tumors with high immune gene expression signatures exhibit improved responses to BRAF-targeted therapy (*45*). Transcriptomic analysis in the COMBI-v cohort further showed the enrichment of gene signatures associated with CD8^+^ T effector cells, cytolytic T cells, antigen presentation and NK cells in complete responder tumors (*11*). Retrospective studies have also reported that high CD8^+^ T cell infiltration is associated with improved overall survival following BRAF-targeted therapy (*15*). In line with these observations, our WES analysis revealed that DR exhibited lower tumor purity compared with RP, suggesting a greater intratumoral immune infiltrate in DR patients. Extending this observation, spatial IHC analysis demonstrated that both the density and spatial localization of CD8^+^ T cells were associated with response to BRAF-targeted therapy. Tumors could be classified as inflamed, excluded, or desert based on CD8^+^ T cell distribution, and only inflamed (high CD8^+^ T cell density within both intratumoral and peritumoral spatial bands) was associated with prolonged PFS. However, our spatial analyses were performed in a limited subset of patient samples due to sample availability, and further validation in independent cohorts will be required before incorporating spatial immune features into predictive models of DR in BRAF-targeted therapy.

Limitations of our study include the fact that we do not have an external validation cohort of patients treated with BRAFi/MEKi in the immunotherapy-naive setting with clinical annotation and inference of intratumoral heterogeneity. However, our results highlight the value of integrating rich clinical data with molecular tumor characterization and the need to generate such multimodal data. Prediction of patients likely to have durable response to targeted therapy may change decision-making and improve outcomes, e.g. in patients who may otherwise be poor candidates for immune checkpoint blockade (e.g. significant autoimmune conditions or comorbid statuses).

In conclusion, our study supports the idea that integrated molecular and clinical features improve prediction of BRAF/MEK targeted therapy response. Genomic heterogeneity integrated with clinical metrics of tumor burden and spatial features of the tumor microenvironment may help identify durable responders who differentially benefit from targeted therapies in BRAF-mutant melanoma.

## Supporting information

Supplementary Table 1 COMBI-d cohort clinical and genomic data

Supplementary Table 2 Recurrent genomic alterations for BRAF-mutant melanoma

Supplementary Table 3 Gain of function alterations in comparison between DR and RP

Supplementary Table 4 Loss of function alterations in comparison between DR and RP

Supplementary Table 5 Spatial CD8+ T-cell distribution and immune phenotype classification in melanoma tumors

## ACKNOWLEDGMENTS

We thank Jasmine Mu and Xiaohong Li for helpful discussions and technical assistance.

## AUTHOR CONTRIBUTION STATEMENTS

Conceptualization: D.L., K.T.F., Y.X.S., A.S., J.C.B., L.A.G., E.V.A.

Methodology: D.L., Y.X.S., B.R., A.S., C.A.R., J.P., G.T., M.P.M., A.H., J.C.B., E.V.A.

Investigation: D.L., K.T.F., D.R., C.A.

Visualization: D.L., K.T.F., Y.X.S., A.S., B.R.

Funding acquisition: D.L., K.T.F., L.A.G.

Project administration: D.L., K.T.F.

Supervision: D.L., K.T.F.

Writing – original draft: D.L., Y.X.S.

Writing – review & editing: D.L., Y.X.S., K.T.F., A.S., B.R., C.A.R., J.P., G.T., M.P.M., A.Y.H., E.V.A., L.A.G.

All authors contributed to the study design, data interpretation, and manuscript preparation, approved the final version of the manuscript, and made the decision to submit it for publication. The authors affirm the accuracy and completeness of the data and adherence of the trial to the protocol.

## MATERIALS AVAILABILITY

Further information and requests for resources and reagents should be directed to and will be fulfilled by DL as corresponding author.

## CODE AVAILABILITY

Software environments and code to reproduce all analyses and generate figures is available on Github (https://github.com/davidliu-lab). Additional reasonable requests for code will be promptly reviewed by the senior authors to verify whether the request is subject to any intellectual property or confidentiality obligations and will be shared to the extent permissible by these obligations.

## DATA AVAILABILITY

All data necessary for figure generation are provided in the Supplementary Tables or are available on GitHub (https://github.com/davidliu-lab). Raw sequencing data will be available in dbGaP (accession number TBD). Any reasonable requests for raw and analyzed data and materials will be promptly reviewed by the senior authors to determine whether the request is subject to any intellectual property or confidentiality obligations. Patient-related information not included in this paper may be subject to patient confidentiality restrictions.

## MATERIALS AND METHODS

### Study design

COMBI-d was a randomized, double-blind study comparing the combination of dabrafenib, a BRAF inhibitor, and trametinib, a MEK inhibitor, with dabrafenib and placebo as first-line therapy in subjects with unresectable (Stage IIIC) or metastatic (Stage IV) BRAF V600E/K mutation-positive cutaneous melanoma. A total of 947 patients were screened for eligibility at 113 participating sites in 14 countries. A total of 423 patients were computer-randomized (1:1) to receive the combination of dabrafenib (150 mg orally twice daily) and trametinib (2 mg orally once daily), or dabrafenib and placebo. Full eligibility criteria have been reported previously (*4*). The primary endpoint of this trial was progression-free survival and overall survival was a secondary endpoint.

### Trial Oversight

The trial was originally sponsored by GlaxoSmithKline. On March 2^nd^, 2015, dabrafenib and trametinib were designated as assets of Novartis, after which Novartis assumed sponsorship of the trial. The study was conducted in accordance with the Declaration of Helsinki and Good Clinical Practice guidelines. The protocol was approved by the institutional review board or independent ethics committee at each participating center. All patients provided written informed consent for data collection supporting these analyses.

### Tissue and Blood Samples

Tumor tissue samples were submitted for central *BRAF* mutation testing at the time of screening. When patient consent permitted, remnant tumor tissue was used for WES and IHC analyses.

Matched germline DNA was used as a reference to identify somatic mutations. FFPE tissue blocks were sectioned at 5 µm thickness. A board certified pathologist reviewed H&E-stained slides to estimate tumor cellularity within the region of interest (ROI) and to measure total area (mm^2^). Based on tumor content, 10 sections were macrodissected and used for DNA extraction.

### WES sequencing

DNA extraction, whole exome library preparation, and sequencing were performed for samples as previously described (*16, 46*). Slides were cut from FFPE blocks and macro-dissected for tumor-enriched tissue. Paraffin was removed from FFPE sections and cores using CitriSolv^TM^ (Fisher Scientific), followed by ethanol washes and tissue lysis overnight at 56°C. Samples were then incubated at 90°C to remove DNA crosslinks, and DNA**–**and when possible, RNA**–**extraction was performed using Qiagen AllPrep DNA/RNA Mini Kit (#51306). Germline DNA was obtained from PBMCs and adjacent normal tissue.

Whole exome capture libraries were constructed from 100 ng of DNA from tumor and normal tissue after sample shearing, end repair, and phosphorylation and ligation to barcoded sequencing adaptors. Ligated DNA was size selected for lengths between 200**–**350 bp and subjected to exonic hybrid capture using Illumina library preparation kits. The samples were multiplexed and sequenced using Illumina HiSeq technology. The Illumina exome uses Illumina’s in-solution DNA probe based hybrid selection method that uses similar principles as the Broad Institute-Agilent Technologies developed in-solution RNA probe based hybrid selection method (*47, 48*) to generate Illumina exome sequencing libraries.

Pooled libraries were normalized to 2 nM and denatured using 0.2 N NaOH prior to sequencing. Flowcell cluster amplification and sequencing were performed according to the manufacturer’s protocols using either the HiSeq 2000 v3 or HiSeq 2500. Each run was a 76-bp paired-end with a dual eight-base index barcode read. Data were analyzed using the Broad Picard Pipeline which includes de-multiplexing and data aggregation.

### Quality control and variant calling

Exome sequence data were analyzed using the Broad Institute CGA pipeline (*49–59*) on the Terra platform, applying the same quality control criteria as previously described (*23, 60*). Specifically, samples were required to meet the following quality control thresholds: mean target coverage > 50× for tumor samples and > 30× for matched normal samples; cross-sample contamination estimation (ContEst) < 5%; tumor purity ≥ 10%, and DeTiN-estimated tumor-in-normal (TiN) ≤ 20%. A power-based filter integrating coverage and tumor purity was applied as described (e.g., minimum 80% power to detect clonal mutations) in (*23, 60*). Seven samples were excluded due to low purity.

MuTect2 (RRID:SCR_026692) (*49*) was used to identify somatic single nucleotide variants (SNVs) in targeted exons, with computational filtering to remove artifacts arising from DNA oxidation during sequencing or FFPE-based DNA extraction using a filter-based method (*52*). Subsequently, Strelka (RRID:SCR_005109) (*51*) was used to identify small insertions or deletions. Finally, Oncotator (RRID:SCR_005183) (*59*) was used to annotate the identified alterations.

### Ploidy and Ploidy Estimation

Purity and ploidy were estimated using the ABSOLUTE algorithm (RRID: SCR_005198) (*56*), which integrates variant allele frequency distributions and copy number variants to estimate absolute tumor purity and ploidy and infer cancer cell fraction (CCF), the proportion of cancer cells in the sample that contain each mutation. Post-purity and ploidy corrected allelic segments were used to estimate allelic copy number estimates.

### Heterogeneity and Aneuploidy estimation

Heterogeneity was estimated here as the proportion of mutations in each sample that were inferred to be subclonal. Clonal mutations were defined as having CCF ≥ 0.8 (*16*), and subclonal mutations were defined otherwise. This definition was selected as a simple, conservative approach with high specificity. To estimate aneuploidy, we used the proportion of the genome inferred to have an allelic amplification or deletion, using allelic segmentation described above.

### Mutational Signatures Calculation

De novo mutational signatures were generated in this cohort using an adaptation of non-negative matrix factorization (*61*) via the Brunet update method (*62*), as previously described in detail (*63*), with the R package SomaticSignatures (RRID:SCR_025620) (*64*) and non-negative matrix factorization (RRID:SCR_023124) (*65*). Cosine similarity was used to compare the discovered signatures to the 30 existing discovered and validated signatures in COSMIC (RRID:SCR_002260) (*66, 67*), with a threshold of 0.85, and we also manually visualized and inspected similarities in mutational motifs between our signatures and COSMIC signatures.

### Functional alteration calling from CNA and mutation data

Functional gene alterations were identified by integrating copy-number alterations, mutation annotations, and OncoKB-supported activating events. CNA and mutation matrices (genes × samples) were harmonized by intersecting samples and taking the union of genes, with missing entries filled as neutral (0).

Somatic variants were first mapped to functional consequence categories. Specifically, protein-truncating alterations-including nonsense mutations, frameshift insertion/deletions, splice-site mutations, start-codon disruptions, and related out-of-frame events were grouped and summarized under a single category, TRUNCATED MUTATION. Missense, in-frame indels, and silent variants were retained as separate categories.

Each gene-sample pair was labeled as gain-of-function (GOF), loss-of-function (LOF), or wild-type (WT). GOF was called if the CNA state indicated amplification (e.g., AMP, HIGH_AMP) or if the gene–sample pair was annotated as activating/hotspot in OncoKB (RRID:SCR_014782) (*25*). LOF was assigned if the CNA state indicated homozygous deletions, if multiple independent mutations were present, or if a two-hit pattern was observed (e.g., LOH together with a TRUNCATED_MUTATION). When both GOF and LOF criteria were satisfied, LOF was given precedence.

### Development and evaluation of a parsimonious predictive model

We developed supervised classification models to identify minimal features predictive of response to BRAF-targeted therapy in the COMBI-d cohort. For DR predictions, patients were classified as DR versus NDR. Tumor genomic heterogeneity and sum of lesion dimensions were modeled as continuous variables, while baseline LDH (normal versus abnormal) and number of metastatic sites (<3 versus ≥3) were treated as categorical variables. Parsimonious non-linear decision tree models were trained using the *rpart* package (method = “class”, RRID:SCR_021777). To prioritize identification of DR, we implemented a cost-sensitive classification scheme by specifying a misclassification loss matrix. False-positive predictions (predicting DR when true class was NDR) were penalized with a cost of 2, whereas false-negative predictions (predicting NDR when the true class was DR) were penalized with a cost of 1. Tree complexity was constrained by limiting the maximum tree depth to 2. Model predictions were generated for all COMBI-d cohort patients and compared with observed outcomes to compute confusion metrics and performance metrics, including sensitivity, specificity, recall, F1 score, accuracy, and Cohen’s kappa. Odds ratios were estimated using Wald confidence intervals. To assess treatment specificity, the decision tree trained on tumor heterogeneity and baseline LDH in the COMBI-d cohort was applied without retraining to an independent, harmonized anti–PD-1-treated metastatic melanoma cohort, including a *BRAF*-mutant subset without prior BRAF-targeted therapy. Model performance was evaluated using the same metrics and outcome definitions across cohorts.

### Survival analysis

Survival analyses were performed utilizing the Python lifelines package (RRID:SCR_024899). For Kaplan–Meier curve survival analysis, a log-rank test (*logrank_test()* function) was used to compare survival curves.

### Hierarchical clustering

Hierarchical clustering was performed using the *clustermap()* function from the Python seaborn package (RRID:SCR_018132), with default settings including a Euclidean distance metric and “single” linkage method of calculating cluster distance (minimization of the nearest point between clusters).

### Tumor infiltration lymphocytes – CD8^+^ T cells assessed by digital pathology

#### Tissue staining

Formalin-fixed paraffin-embedded (FFPE) tissue sections (4–5 µm thickness) from a subset of pretreatment metastatic melanoma samples were subjected to dual-plex immunohistochemistry (IHC). Staining was performed on the VENTANA BenchMark ULTRA platform using the ultraView^TM^ Universal DAB detection kit and ultraView^TM^ Universal AP Red detection kit (VENTANA).

Sections were stained with hematoxylin, a melanoma triple cocktail (MTC; HMB45, A103, and T311), and an anti-CD8 antibody (rabbit monoclonal, clone SP57). Both CD8 and MTC antibodies were pre-diluted, ready-to-use antibodies from VENTANA. The MTC was used to identify melanocytic tumor cells, while CD8 staining identified infiltrating cytotoxic T cells.

Assay performance and reproducibility were validated through intra-run precision testing across multiple staining runs. Each run included appropriate duplex negative controls (mouse and rabbit IgG) and a run control sample, with no detectable staining observed in negative controls. All staining procedures were conducted according to validated protocols established by the external vendor.

#### Digital imaging acquisition and HALO analysis

A dualplex immunohistochemistry (IHC) assay targeting CD8 and MTC was developed. Stained slides were scanned at 20× magnification using an Aperio AT2 Leica digital whole-slide scanner. Digital image analysis was performed using the multiplex IHC module (v1.2) within the HALO software platform (v2.2; Indica Labs). Lymphoid tissues were excluded from analysis.

Melanoma lesions were annotated by a pathologist in close collaboration with imaging scientists. Tumor regions and invasive margins were annotated based on tumor cell morphology, with MTC staining serving as a reference. The invasive margin was defined as the interface between tumor (MTC^+^ region) and adjacent stromal tissue, guided by both MTC and hematoxylin staining.

A customized image analysis algorithm was developed to quantitatively assess the percentage of CD8-positive cells within tumor and stromal compartments. To evaluate the spatial distribution of CD8^+^ T cells (infiltration analysis), a peritumoral region was defined as extending 150 µm on either side of the tumor–stromal boundary. This region was further subdivided into concentric distance bins at 30 µm intervals, spanning both intratumoral and stromal compartments. CD8^+^ T cell density was quantified within each bin by normalizing CD8^+^ T cell counts to the corresponding area, yielding cell density (cells/mm²) to generate spatial distribution profiles. For summary analysis, intratumoral CD8^+^ T cell density was defined as the mean density across intratumoral bands (–150 to –30 μm; bands –5 to –1), while peritumoral CD8^+^ T cell density was defined as the mean density across stromal bands (30 to 150 μm; bands 1 to 5).

## Statistics and reproducibility

Statistical tests were performed utilizing the Python *scipy.stats* package (RRID:SCR_008058). To compare numeric features between response categories, a non-parametric Mann–Whitney–Wilcoxon (MWW) rank-sum test (*mannwhitneyu()* function) was used to minimize the effects of outliers. For comparison of proportions across response categories, a chi-squared test (*chi2_contingency()* function) was utilized. For associations between binary variables (e.g. association of gene alteration with responders vs. progressors), a Fisher’s exact Test (*fisher_exact()* function) was utilized to generate a *P* value. A conservative adjusted odds ratio (OR) was generated by repeating the Fisher’s Exact test adding one to both the number of gene mutant responders and progressors. All tests were two-sided unless otherwise indicated.

