## Supplementary Figures for "Genomic, Clinical, and Spatial Predictors of Durable Response to BRAF/MEK Inhibition in *BRAF*-Mutant Melanoma"

Supplementary Figure 1.

**A.**

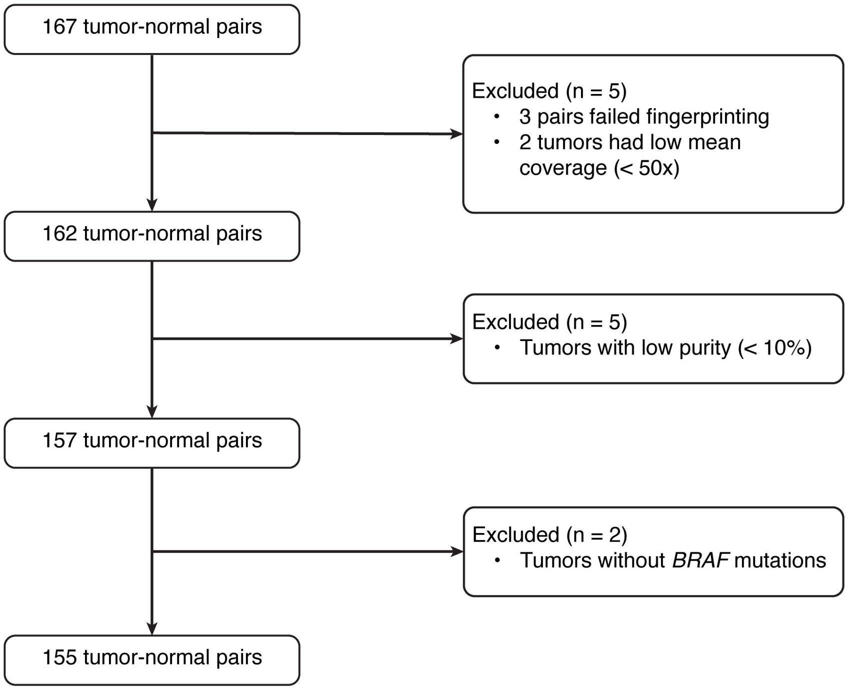

**Supplementary Figure 1. COMB-d cohort sample QC process. (A)** Consort diagram showing inclusion, exclusion, and quality control criteria for patients/tumors included in the analysis.

Supplementary Figure 2.

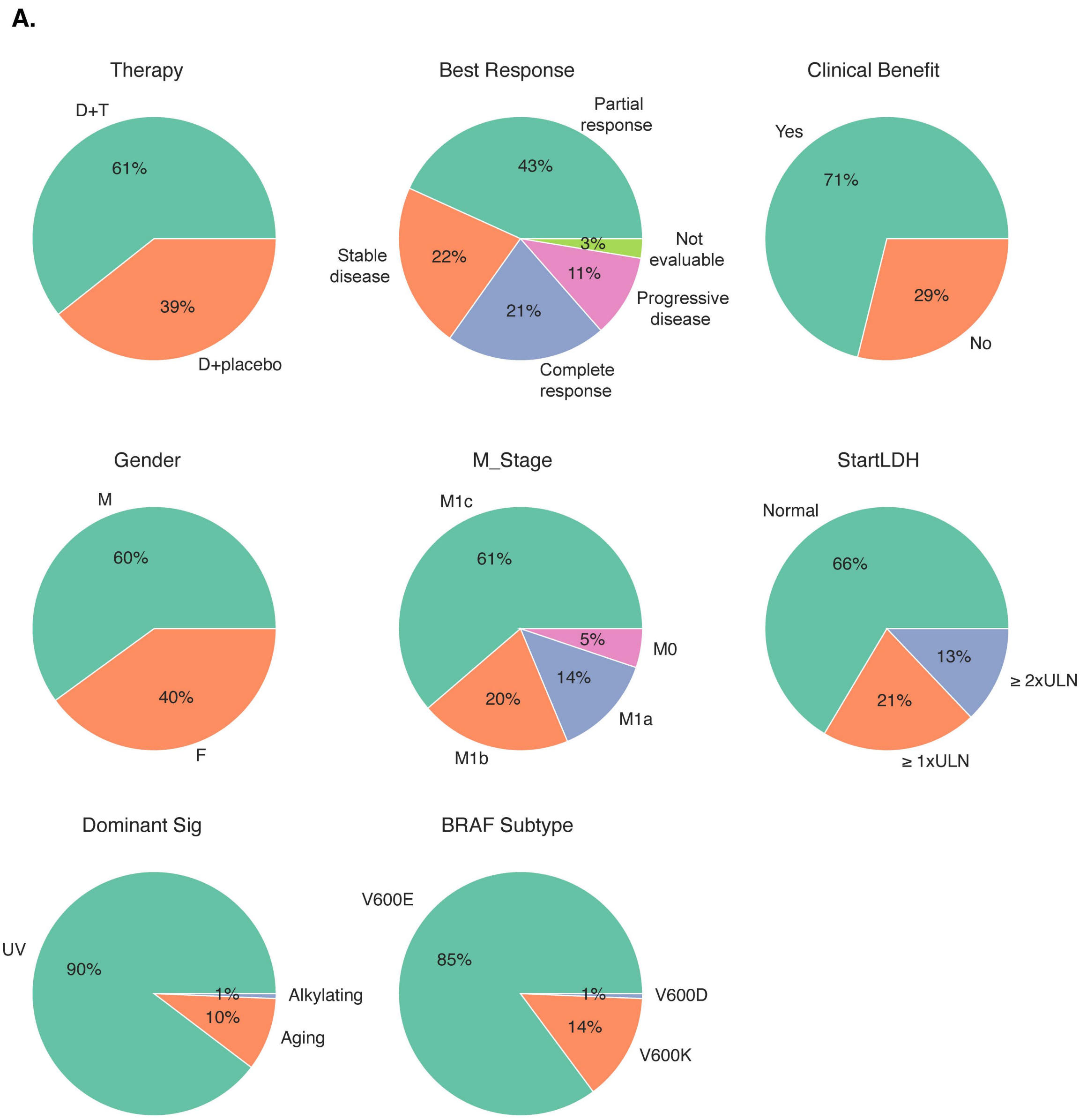

**Supplementary Figure 2. Clinical and genomic feature distribution of the COMBI-d cohort.** Summary of treatment, response, clinical and genomic features of patients in the COMBI-d cohort.

### Supplementary Figure 3.

**A.**

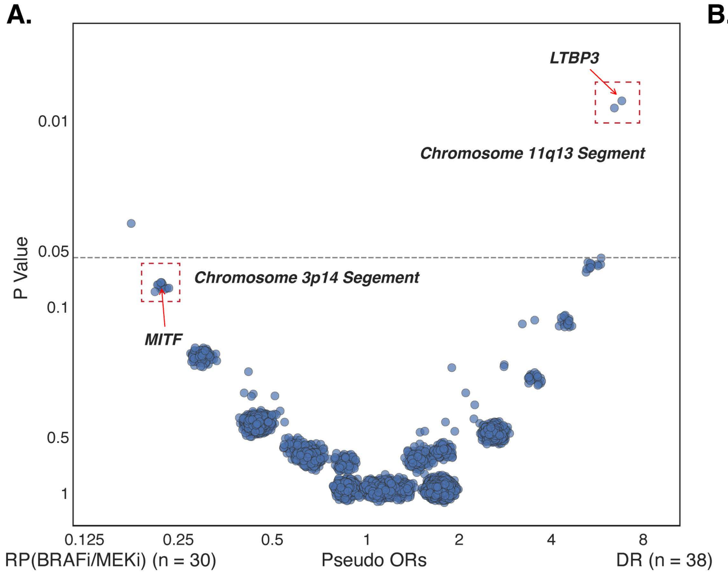


**B.**

| *Chr11q13 (LTBP3)* | Responders (DR) | Nonresponders (RP. BRAFi/MEKi) | Intermediate Group (INT) | Total |
|---|---|---|---|---|
| Amp | 11 | 1 | 5 | 17 |
| No Amp | 27 | 29 | 82 | 138 |
| | 38 | 30 | 87 | 155 |
| *Chr3p14 (MITF)* | Responders (DR) | Nonresponders (RP. BRAFi/MEKi) | Intermediate Group (INT) | Total |
| Amp | 0 | 3 | 2 (progressed after 1 year) | 5 |
| No Amp | 38 | 27 | 85 | 150 |
| | 38 | 30 | 87 | 155 |

**C.** **D.** **E.** **F.**

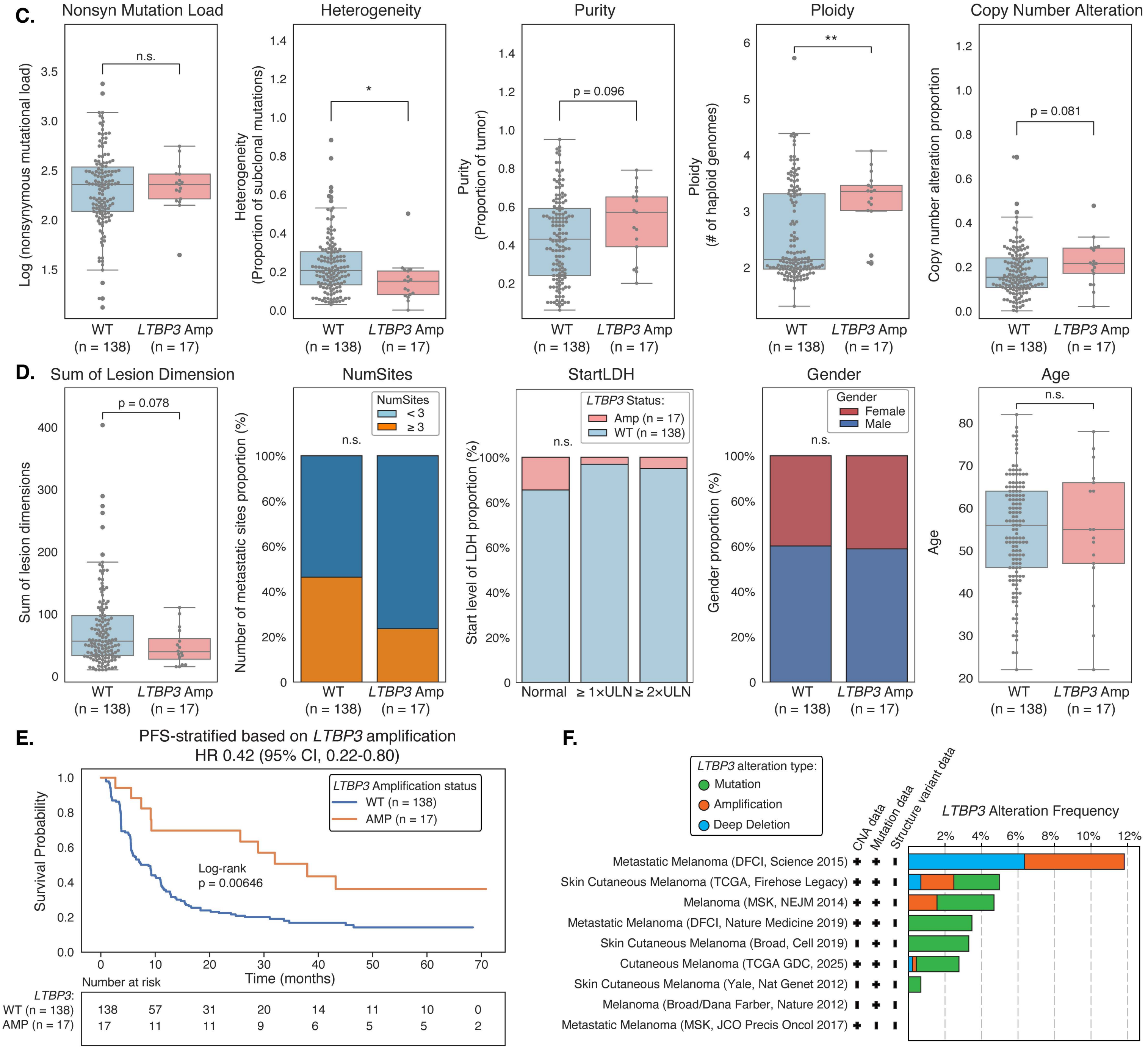

**Supplementary Figure 3. *LTBP3* copy number amplification in the COMBI-d cohort and external melanoma cohorts.** (**A**) Comparison of gain of function alteration frequencies between RP and DR. One patient was excluded because copy number alterations could not be assessed due to low tumor purity. Group comparisons were performed using Fisher's exact test. (**B**) Contingency table showing the distribution of DR, RP, and INT by copy number amplification status at *Chr11q13 (LTBP3)* and *Chr3p14 (MITF)*. (**C**) Boxplots showing the comparison of genomic features between tumors with (n = 17) and without (n = 138) *LTBP3* amplification, including log-transformed nonsynonymous mutational load, genomic heterogeneity, tumor purity, ploidy, and the proportion of copy number alterations. Group comparisons were performed using two-sided Mann–Whitney–Wilcoxon tests with Bonferroni correction; corresponding P values are shown. (**D**) Boxplots and stacked bar plots showing the comparison of clinical features between tumors with and without *LTBP3* amplification, including sum of lesion dimensions, number of metastatic sites, baseline LDH level, sex and age. Continuous variables were compared using two-sided Mann-Whitney-Wilcoxon tests with Bonferroni correction, and categorical variables using chi-squared tests; corresponding P values are shown. (**E**) Kaplan–Meier PFS curves stratified by *LTBP3* amplification status. P values were calculated using the log-rank test. (**F**) Frequency of *LTBP3* amplification across selected melanoma cohorts from TCGA with available copy number alteration data, including DFCI (Science 2015), TCGA (Firehose Legacy), MSK (NEJM 2014), DFCI (Nature Medicine 2019), Broad (Cell 2019), TCGA GDC (2025), Yale (Nature Genetics 2012), Broad/DFCI (Nature 2012), MSK (JCO Precis Oncol 2017).

### Supplementary Figure 4.

**A.**

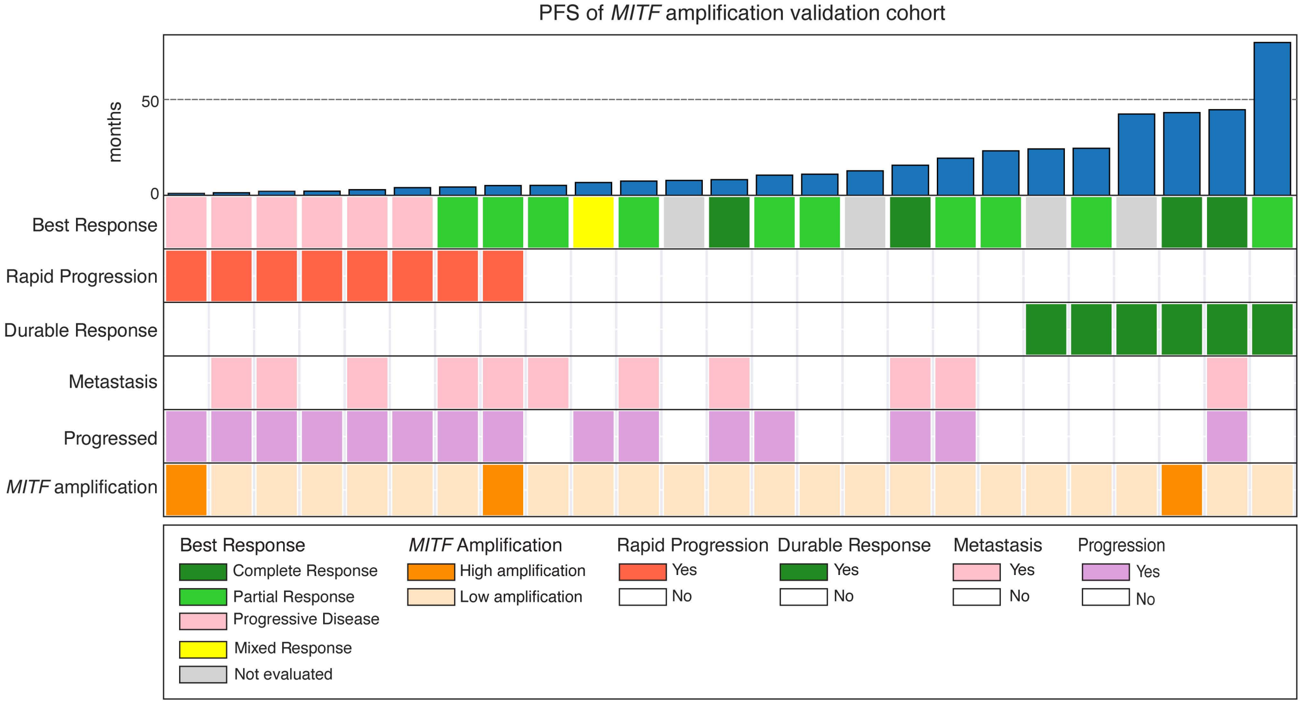


**B.**

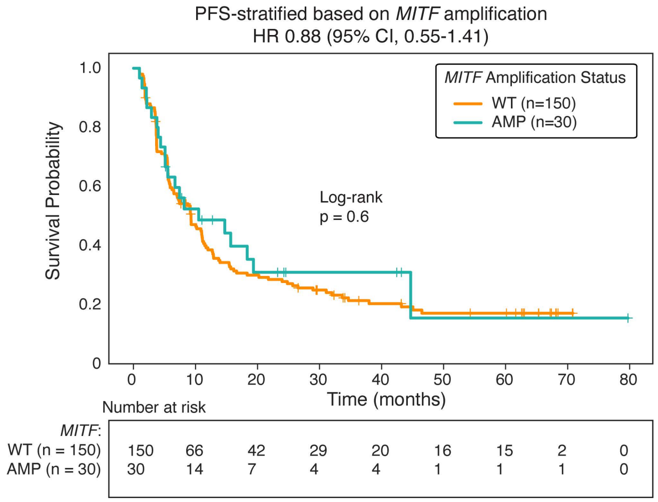


**C.**

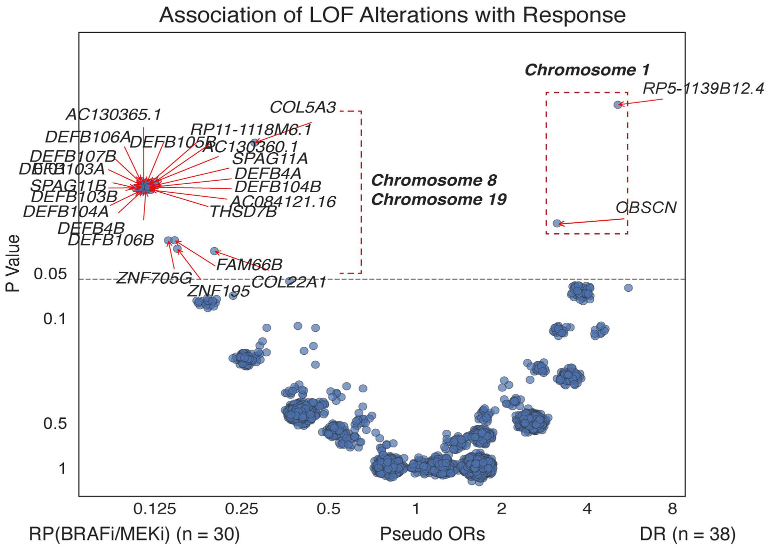

**Supplementary Figure 4. Validation of *MITF* amplification associated with RP in BRAF-targeted therapy.** (**A**) CoMut plot showing clinical and genomic characteristics in the *MITF* amplification validation cohort. Each column represents a tumor, ordered by PFS in ascending order. Best response was defined according to RECIST criteria: CR (green), PR (light green), PD (pink), mixed response (MS; yellow), and not evaluated (grey). Disease stage is indicated, with metastatic melanoma shown in pink. *MITF* amplification status (high versus low) was defined based on copy number estimates from the DFCI Oncopanel. (**B**) Kaplan–Meier PFS curves stratified by *MITF* amplification status in the combined cohort (COMBI-d and MITF validation cohorts). P values were calculated using the log-rank test. (**C**) Comparison of LOF alteration frequencies between RP and DR. One patient was excluded because copy number alterations could not be assessed due to low tumor purity. Red dashed boxes highlight clusters of genes located on chromosomes 1, 8 and 19 that are associated with response. Group comparisons were performed using Fisher's exact test.

Supplementary Figure 5.

A.

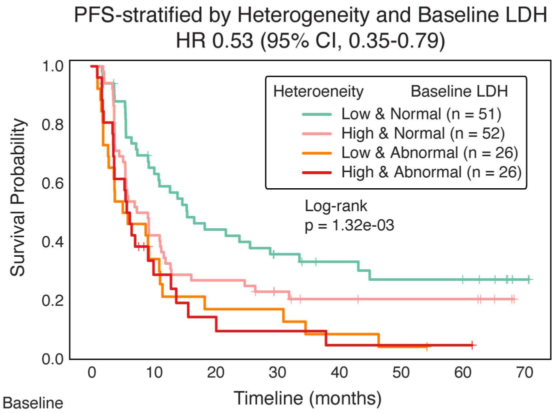


| Heteroeneity Baseline LDH | Number at risk | | | | | | | |
|---|---|---|---|---|---|---|---|---|
| Low & Normal (n = 51) | 51 | 31 | 21 | 14 | 11 | 9 | 9 | 2 |
| High & Normal (n = 52) | 52 | 22 | 14 | 9 | 6 | 5 | 5 | 0 |
| Low & Abnormal (n = 26) | 26 | 8 | 4 | 4 | 2 | 1 | 0 | 0 |
| High & Abnormal (n = 26) | 26 | 7 | 3 | 2 | 1 | 1 | 1 | 0 |

B.

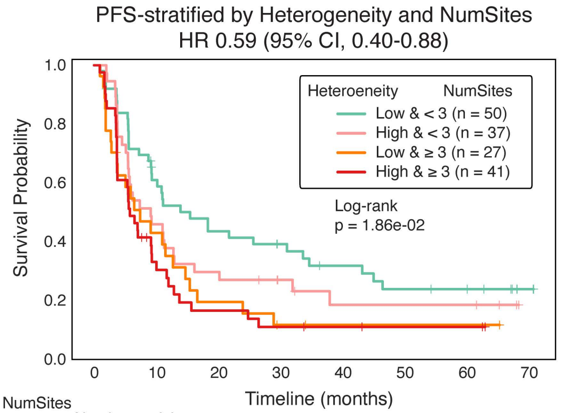


| Heteroeneity NumSites | Number at risk | | | | | | | |
|---|---|---|---|---|---|---|---|---|
| Low & < 3 (n = 50) | 50 | 28 | 20 | 16 | 12 | 9 | 8 | 2 |
| High & < 3 (n = 37) | 37 | 17 | 11 | 7 | 4 | 4 | 4 | 0 |
| Low & ≥ 3(n = 27) | 27 | 11 | 5 | 2 | 1 | 1 | 1 | 0 |
| High & ≥ 3 (n = 41) | 41 | 12 | 6 | 4 | 3 | 2 | 2 | 0 |

C.

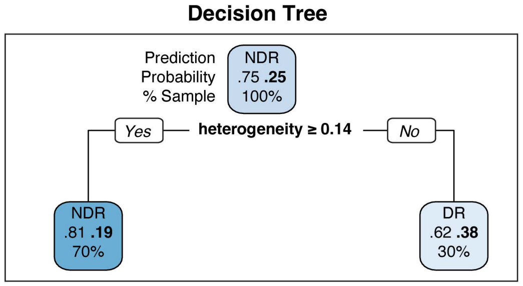


D.

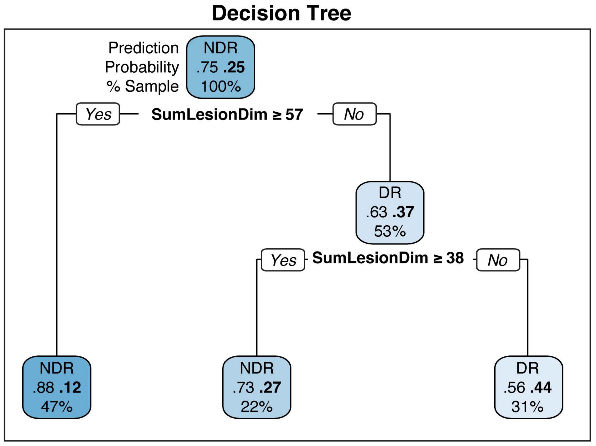


E.

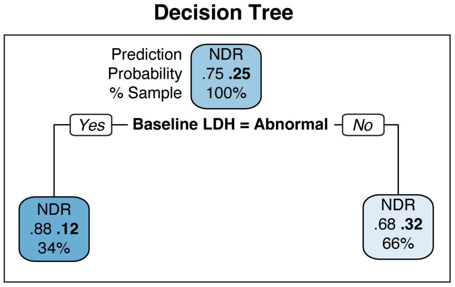


F.

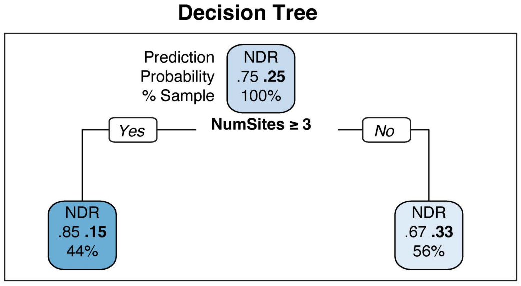

**Supplementary Figure 5. Decision tree models using genomic heterogeneity and tumor burden metrics alone to predict DR in BRAF-targeted therapy.** (**A**) Kaplan–Meier PFS curves stratifying patients by median dichotomization of genomic heterogeneity and baseline LDH (normal versus abnormal) into four groups. P values were calculated using the log-rank test. (**B**) Kaplan–Meier PFS curves stratifying patients by median dichotomization of genomic heterogeneity and number of metastatic sites (<3 versus ≥3) into four groups. P values were calculated using the log-rank test. (**C–F**) Structure of the decision tree model used to classify DR versus NDR based on genomic heterogeneity (**C**), sum of lesion dimensions (**D**), baseline LDH (**E**), and number of metastatic sites (**F**). For each node, the predicted class is shown at the top, the estimated probabilities of NDR (left) and DR (right) are shown in the middle, and the percentage of patients assigned to the node is shown at the bottom.

Supplementary Figure 6.

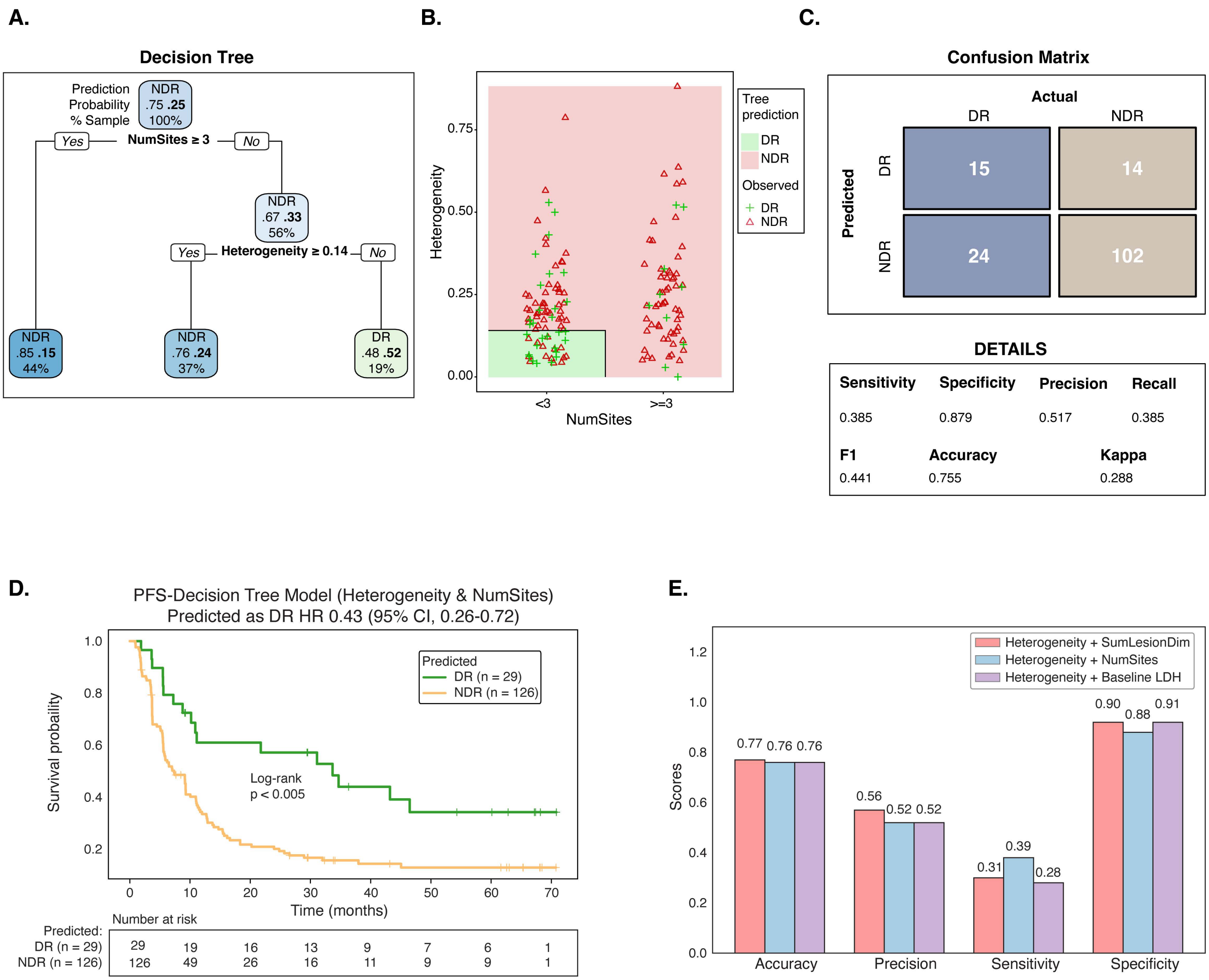

**Supplementary Figure 6. Decision tree models combining genomic heterogeneity with individual tumor burden metrics to predict DR in BRAF-targeted therapy.** (**A**) Structure of the decision tree model used to classify DR versus NDR based on tumor genomic heterogeneity and number of metastatic sites. For each node, the predicted class is shown at the top, the estimated probabilities of NDR (left) and DR (right) are shown in the middle, and the percentage of patients assigned to the node is shown at the bottom. (**B**) Decision boundaries defined by thresholds on tumor genomic heterogeneity and number of metastatic sites. Background shading indicates the model-predicted probability of DR, where $p = P(DR)$; $p \geq 0.5$ is classified as DR and $p < 0.5$ as NDR. Individual patients are overlaid, with green crosses denoting true DR and red triangles denoting true NDR. (**C**) Confusion matrix summarizing model performance in the cohort, showing predicted versus observed response categories. (**D**) Kaplan–Meier PFS curves stratified by decision tree–predicted DR and NDR groups in the COMBI-d cohort. P values were calculated using the log-rank test. (**E**) Bar plots showing prediction performance metrics for decision tree models trained to predict DR using tumor heterogeneity combined with different tumor burden-associated clinical features: sum of lesion dimensions, number of metastatic sites, and baseline LDH. Performance metrics shown include accuracy, precision, sensitivity, and specificity for each model.

### Supplementary Figure 7.

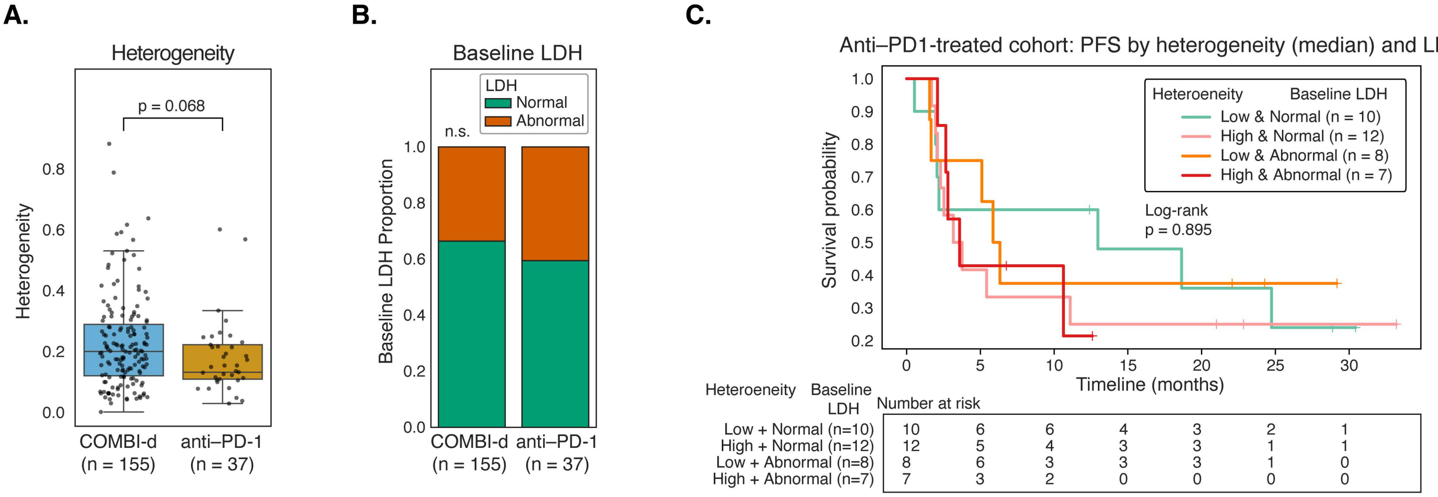

**Supplementary Figure 7. Tumor genomic heterogeneity and baseline LDH in the Anti–PD-1 cohort.** (**A**) Boxplots comparing baseline tumor genomic heterogeneity between the COMBI-d cohort and the harmonized anti-PD1 cohort. Group comparisons were performed using a two-sided Mann–Whitney–Wilcoxon test; the corresponding P value is shown. (**B**) Proportion of patients with normal versus abnormal baseline LDH levels in the COMBI-d and anti–PD-1 cohorts. Group differences were assessed using a chi-squared test. (**C**) Kaplan–Meier progression-free survival (PFS) curves for the anti–PD-1 cohort stratified by median dichotomization of tumor heterogeneity (high vs low) and baseline LDH status (normal vs abnormal). P values were calculated using the log-rank test. The number of patients at risk at each time point is shown below.

Supplementary Figure 8.

A.

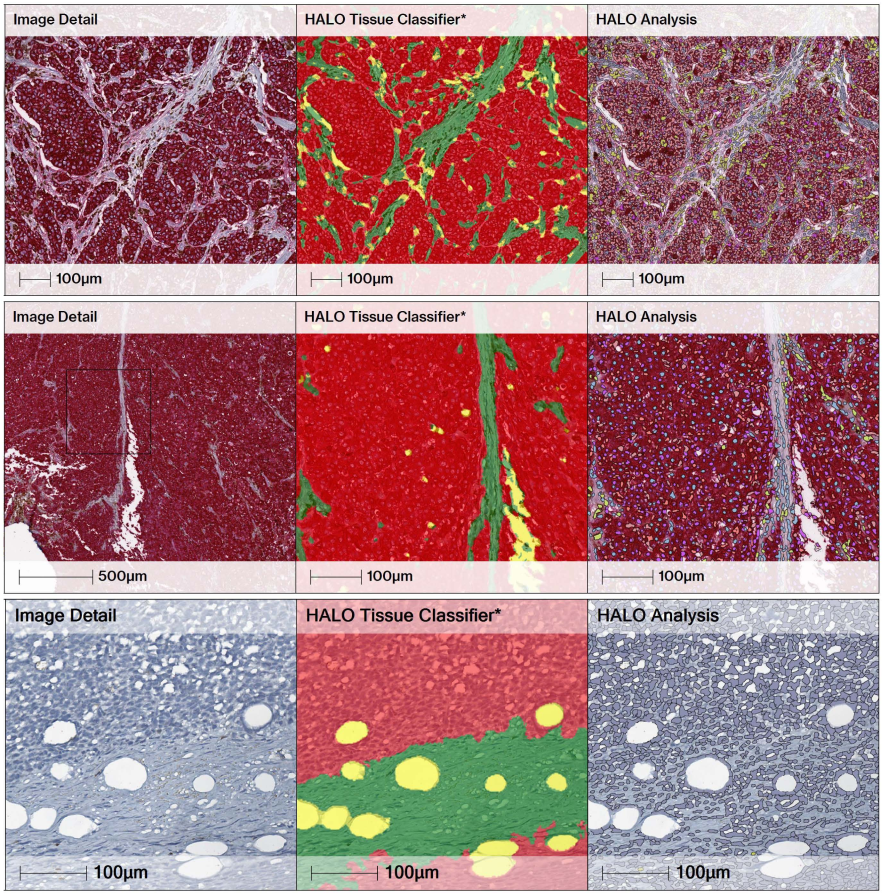


HALO Analysis Phenotype Legend:

- Hematoxylin$^{+}$ cells
- CD8$^{+}$ T cells
- Hemtoxylin$^{+}$, MTC$^{+}$ T cells
- MTC$^{+}$ cells
- CD8$^{+}$, MTC$^{+}$ cells
- Hematoxylin$^{+}$, CD8$^{+}$, MTC$^{+}$ cells

HALO Analysis Tissue Classification:

Tumor: Red
Stroma: Green
Excluded: Yellow

B.

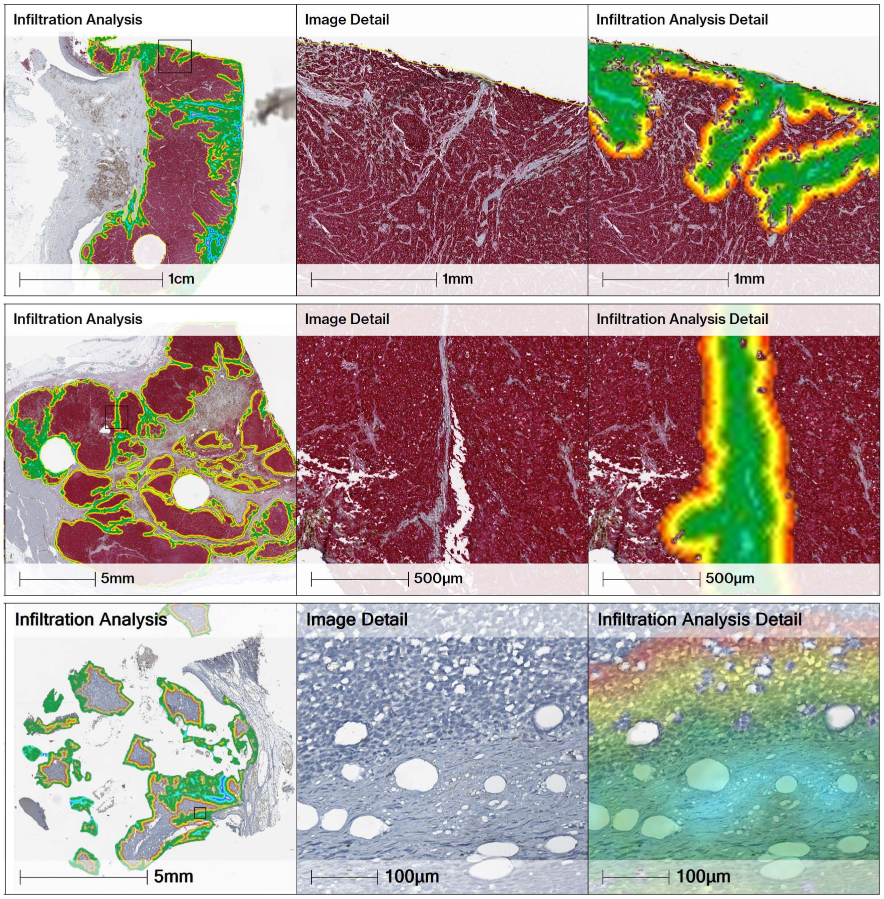

**Supplementary Figure 8. HALO-based tissue classifier, cell-level phenotype analysis, and infiltration analysis of CD8$^+$ T cell infiltration in the COMBI-d cohort.** (**A**) Representative images illustrating tissue segmentation and cell phenotyping workflows used for spatial proximity analysis of CD8$^+$ T cells. *Column 1*: Higher-magnification IHC images corresponding to Fig. 5A (*Column 1*), showing melanoma tumor lesions and adjacent stromal regions. Scale bars, 100 µm and 500 µm. *Column 2*: A tissue segmentation classifier, developed and validated using HALO Tissue Classifier Analysis, distinguishes tumor and stromal compartments by recognizing hematoxylin and MTC staining. Tumor (red) and stroma (green) regions are identified based on morphological and cellular features of MTC-stained melanoma cells, with excluded areas (yellow) marking empty glass slides, background, or necrosis. Scale bars, 100 µm. *Column 3*: HALO image analysis groups cells based on their shape and marker expression, identifying cell types such as CD8$^+$ T cells and MTC$^+$ tumor cells. Colored overlays display various IHC marker combinations for cell phenotypes, with details provided in the legend. Scale bars, 100 µm. (**B**) Representative images of HALO-based CD8$^+$ T cell infiltration analysis across inflamed, excluded, and desert tumor phenotypes. *Column 1*: Whole-slide MART1 IHC sections analyzed using HALO digital pathology software to identify tumor regions and quantify CD8$^+$ T cell proximity to the tumor boundary. Black rectangle boxes denote magnified regions shown in Columns 2 and 3. Scale bars, 1 cm and 5 mm. *Column 2*: Magnified histologic views of the regions indicated in *Column 1*, showing the tumor–stromal interface. Scale bars, 1 mm, 500 µm, and 100 µm. *Column 3*: Corresponding HALO infiltration heatmaps illustrating the spatial gradient of CD8$^+$ T cell proximity relative to the tumor boundary, with warmer colors indicating regions closer to the tumor margin and cooler colors representing regions farther from the tumor interface. Scale bars, 1 mm, 500 µm, and 100 µm.

Supplementary Figure 9.

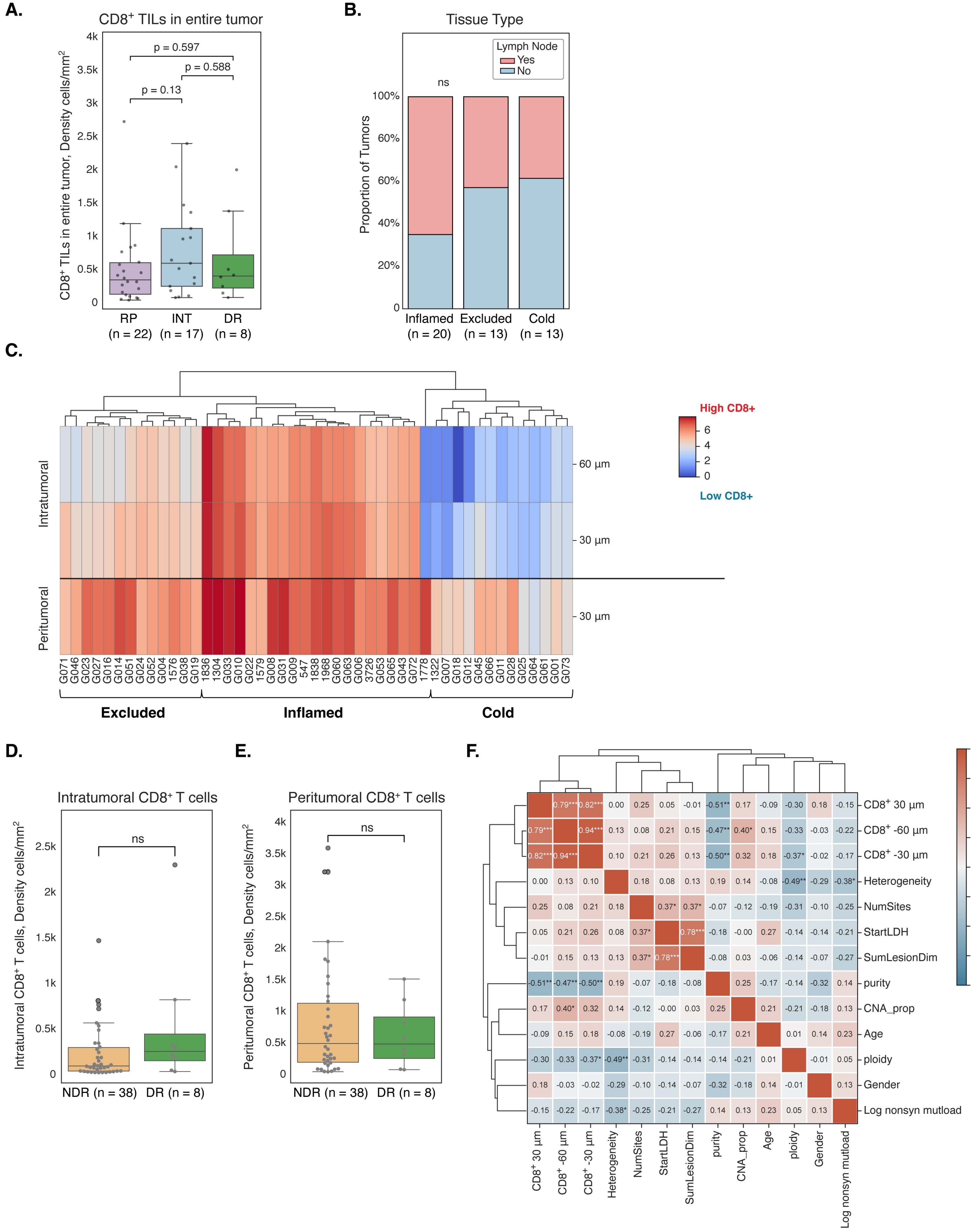

**Supplementary Figure 9. CD8$^+$ T cell infiltration quantification in the COMBI-d cohort.** (**A**) Boxplot showing the comparison of CD8$^+$ T cell proportion across the entire tumor between RP, INT, and DR. Group comparisons were performed using a two-sided MWW test with Bonferroni correction; the corresponding P value is shown. (**B**) Stacked bar plot showing the proportion of tumor samples collected from lymph nodes (pink) across inflamed, excluded, and cold tumor immune phenotypes. Group comparisons were performed using a chi-squared test; the corresponding P value is shown. (**C**) Hierarchical clustering of tumors based on CD8$^+$ T cell density in the intratumoral (within 60 µm of the tumor boundary) and peritumoral regions. (**D–E**) Boxplots showing the comparison of intratumoral (**D**) and peritumoral (**E**) CD8$^+$ T cell density between DR and NDR. Intratumoral CD8$^+$ T cells were calculated as the mean density across band –5 to –1, and peritumoral CD8$^+$ T cells as the mean density across bands 1 to 5. Group comparisons were performed using two-sided MWW tests with Bonferroni correction; corresponding P values are shown. (**F**) Heatmap showing pairwise Pearson correlation coefficients among CD8$^+$ T cell densities, genomic and clinical features in the COMBI-d cohort. Features are ordered by hierarchical clustering based on correlation distance. Pearson correlation coefficients are showed within each cell, with asterisk annotation indicating nominal significance levels as shown. Color scale represents the strength and direction of correlation. Asterisks denote nominal significance (*: $P < 0.05$;**: $P < 0.01$;***: $P < 0.001$).
